# Immunoglobulin A Carries Sulfated *N-*glycans Primarily at the Tailpiece Site – An Oxonium-Ion-Guided Approach for Site-Specific *N*-glycan Identification

**DOI:** 10.1101/2024.06.06.597690

**Authors:** Frania J. Zuniga-Banuelos, Greta Lemke, Marcus Hoffmann, Udo Reichl, Erdmann Rapp

## Abstract

Sulfated *N-*glycans from human immunoglobulin A (IgA) were recently discovered via glycomic approaches. However, their site-specific description is still pending. Certain *N-*glycan structures at specific *N-*glycosylation sites in IgA are crucial for microbial neutralization and effector functions. For instance, sialylated *N-*glycans on the C-terminal tailpiece mediate anti-viral activity by interfering with sialic-acid-binding viruses. Sulfated *N-*glycan epitopes can be ligands for viral proteins and thus play a role in the immune response. In this study, we performed a site-specific screening for sulfated *N-*glycans in two commercially available human serum IgA samples employing an in-depth *N-*glycoproteomic approach, previously developed by us. We found evidence of complex-type and hybrid-type *N-*glycans containing sulfated *N-*acetylhexosamine (sulfated HexNAc) attached to the *N-*glycosylation sites in the tailpiece and the C_H_2 domain of both IgA subclasses. A detailed comparison of the *N-*glycosylation profiles of human serum IgA samples from two suppliers showed such *N-*glycans with sulfated HexNAc consistently in higher abundance in the tailpiece region. Surprisingly, also complex-type *N-*glycan compositions bearing *O-*acetylated sialic acid were identified in the tailpiece. These findings have not been described before for a site-specific glycopeptide analysis. Overall, our work provides a methodology for performing a dedicated site-specific search for sulfated and *O-*acetylated *N-*glycans that can be easily transferred, e.g. to human IgA derived from mucosal tissues, milk, or saliva. Our future aim is to include sulfated *N-*glycans into longitudinal studies of IgA *N-*glycosylation and to investigate their role as a biomarker and a treatment option.

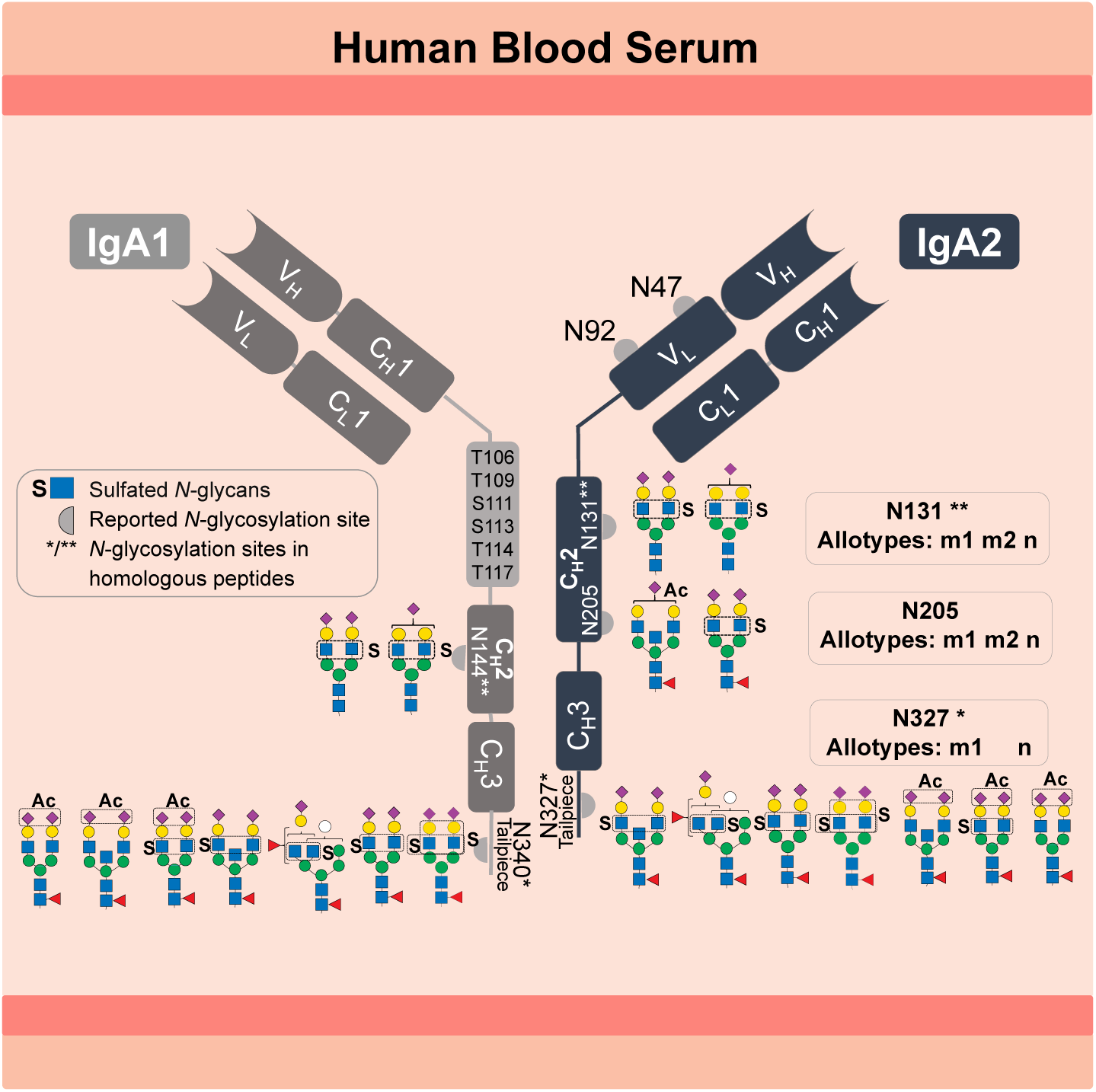

## Introduction

Human immunoglobulin A (IgA) is the most abundant secretory immunoglobulin and the second most abundant in serum (1, 2). IgA plays a crucial role in the defense against mucosal infections, microbiota modulation, and newborn immunization (3–5). Due to its potent antiviral activity, tumor cell killing effector mechanism and ability to inhibit inflammatory and autoimmune diseases, IgA has an enormous therapeutic potential (6–8). In humans, IgA exists in two subclasses, IgA1 and IgA2 (ratio 89 to 11 in serum) (1). Variants with amino acid substitutions exist for both subclasses (e.g. IgA2 M319L related to UniProt P01877 and P0DOX2). IgA2 exists in three genetically controlled polymorphisms (allotypes) m1, m2, and n, whose ratio in serum varies among different ethnical groups (9–11). The highly *O-*glycosylated hinge region of IgA1 is 13 amino acids longer than the IgA2 hinge region (UniProt P01876) (1). IgA1 and IgA2 are present in two structural conformations: as dimer or monomer. In dimeric conformation transport across the epithelial barriers is mediated by binding to the polymeric immunoglobulin receptor (pIgR), the precursor of the secretory component required to form secretory IgA (1). From 87% to 97% of IgA secreted into mucosal tissues (intestinal, nasal, and oral tissues) are dimerized via the joining-chain (J-chain). In contrast, from 85% to 99% of the IgA present in blood serum exist as monomers (2). Both IgA subclasses are modified with *N-*glycans and share homologue sequences around the *N-*glycosylation sites in the C_H_2 domain (N144-IgA1/N131-IgA2_m1/m2/n_), as well as on the C-terminal tailpiece in C_H_3 domain (N340-IgA1/N327-IgA2_m1/n_) – the allotype m2 is not homologous to the m1 and n allotypes in the tailpiece sequence (1, 11). All IgA2 allotypes have two additional *N-*glycosylation sites, one in the C_H_2 domain and another in the C_H_1 domain (N205 and N47) but only the allotypes m2 and n have a fifth *N-*glycosylation site (N92) in the C_H_1 domain (1, 11). For better clarity, the graphical abstract provides an overview of the information presented above.

The *N-*glycosylation profile varies between IgA subclasses and *N-*glycosylation sites. On the one hand, a lectin blot analysis on each serum IgA subclass revealed that IgA2 bears fewer sialylated, galactosylated and bisected *N-*glycans and more hybrid- and oligomannose-type *N-*glycans than IgA1 (12). Furthermore, these specific *N-*glycosylation profiles seem to be critical for the pro-inflammatory effect of IgA2 and the anti-inflammatory modulatory role of IgA1 (12). On the other hand, several studies have used bottom-up mass spectrometry analyses to obtain a site-specific description of the serum IgA *N-*glycosylation (12–17). However, the *N-*glycans observed at the N144-IgA1 and N131-IgA2_m1/m2/n_ sites (in the C_H_2 domain), as well as the tailpiece sites N340-IgA1 and N327-IgA2_m1/n_ (in the C_H_3 domain), are reported to both IgA subclasses due to the homologous sequences surrounding these sites. Yet, significant differences between the *N-*glycosylation profile of both C_H_2 and C_H_3 domains were observed. The C_H_2 domain features mostly non-fucosylated hybrid-type *N-*glycans and di-antennary complex-type *N-*glycans with terminal galactose or sialic acid. In contrast, the tailpiece typically bears fucosylated di- or multi-antennary complex-type *N-*glycans variably sialylated and bisected, as well as oligomannose-type *N-*glycans. Some studies show that *N-*glycans attached to the sites in the C_H_2 domain and the tailpiece (IgA1 Fc *N-*glycans) are irrelevant for binding to the FcαRI receptor (13, 18). However, other studies indicate that IgA1 Fc *N-*glycans are relevant for other receptors involved in anti-inflammatory mechanisms (8, 12). Closer investigations on the *N-*glycosylation at the tailpiece have demonstrated its high impact on different IgA functions. First, *N-*glycosylation at the tailpiece is critical for binding to complement C3 protein (19). Second, it was shown to be important for modulating dimer formation in IgA1 (19). Third, *N-*glycosylation at the tailpiece enhances the IgA anti-viral neutralizing activity by exposing sialylated *N-*glycans and interacting with sialic-acid-binding viral proteins, like influenza hemagglutinin and neuraminidase (6). Fourth, it affects the serum half-life of IgA1 (20). Rifai *et al.* showed that the clearance of IgA1 from serum is slower when *N-*glycans at the tailpiece are removed, whereas it remained unchanged when *O-*glycans are removed (20). Although the function of IgA2 *N-*glycans on the C_H_1 domain (Fab region) remains unclear so far, Rifai *et al.* suggests that the additional *N-*glycosylation sites on IgA2 (N47 and N92 in the C_H_1 and N205 in C_H_2) alter the rate of IgA2 clearance from blood. These findings demonstrate the significance of elucidating protein *N-*glycosylation in terms of micro-heterogeneity (describing the site-specific *N-*glycan variability), for the comprehension of its role in the mechanisms modulating the effector functions of IgA.

In addition to the complexity caused by glycosylation, structural conformations, subclasses, and polymorphisms, IgA was identified to have a prominent abundance of *N-*glycans modified by sulfation. This was detected for the first time in 1999 by Boisgard *et al.* in IgA from mammary glands of rabbit in addition to other species and tissues (21). However, this could be confirmed for human serum IgA only 20 years later by in-depth glycomic analyses (21–23). Identification of sulfated *N-*glycans is a challenging task, as in several other glycomic and glycoproteomic analyses conducted on IgA, sulfated *N-*glycans never appeared in the identification lists (12–17, 24). Recently Cajic *et al*. and Chuzel *et al.* confirmed the presence of the sulfated *N-*glycan FA2G2S2-SO_4_ in human serum IgA (22, 23). Chuzel *et al.* demonstrated that the sulfate was linked to the 6-carbon of GlcNAc (GlcNAc-6-SO_4_) with the application of a highly-specific sulfatase in combination with the methodology developed by Cajic *et al.* (22). In this methodology, the sulfated *N-*glycan FA2G2S2-SO_4_ was released upon cleavage of IgA *N-*glycans and all together were labeled with the removable fluorescent dye 9-fluorenylmethyl chloroformate (Fmoc) (23). The Fmoc-labeled sulfated *N-*glycan was isolated by hydrophilic interaction high-performance liquid chromatography (HILIC-HPLC), identified, and characterized employing two orthogonal approaches: one based on matrix-assisted laser desorption/ionization time-of-flight mass spectrometry (MALDI-TOF-MS) and another based on multiplexed capillary gel electrophoresis with laser-induced fluorescence detection (xCGE-LIF) *N-*glycan analysis (23). Cajic *et al*. proved with these approaches, that the newly discovered sulfated *N-*glycan, which composition is HexNAc_4_Hex_5_Fuc_1_NeuAc_2_Sulfo_1_, presents α2-6-Neu5Ac on both antennae and GlcNAc-6-SO_4_ on the α1-3-Man arm, yet without determining the glycosylation sites (23). Alagesan *et al.* reported other sulfated *N-*glycans bearing sulfated galactose, which were associated with the heavy chains of both human IgA subclasses but not assigned to specific *N-*glycosylation sites (25). These galactose sulfated *N-*glycans were deposited in GlyConnect (26, 27).

Although the role of sulfated *N-*glycans in IgA still remains largely unexplored, it is evident that the role of sulfated glycans in general is not minor, since they impact cell-cell interaction and, as HexNAc-sulfated sialosides, can be ligand for some influenza hemagglutinin variants (28–32). Thus, in spite of their relevance, the site-specific detection of the low-abundant sulfated *N-*glycans by mass spectrometry is still pending due to their even lower abundance per glycosylation site, inadequate data analysis, and limitations associated with mass spectrometry (MS) measurement – the negative charge on sulfated sugars causes low ionization efficiency and instability of the sulfated fragment ions (33).

Therefore, we here determined the *N-*glycosylation sites of human IgA that harbor sulfated *N-*glycans using an in-depth *N-*glycoproteomic approach, with which for the first time, the site-specific identification of HexNAc-sulfated *N-*glycans in human IgA was achieved. The approaches developed by us in previous works allow the identification of intact *N-*glycopeptides from the low-abundant *N-*glycoproteome including *N-*glycopeptides that feature sulfated, glucuronidated or *O-*acetylated *N-*glycans (34, 35). The present study reveals that sulfated *N-*glycans are linked to the C_H_2 domain sites N205, N131 in IgA2 and N144 in IgA1, as well as in the tailpiece site N340 in IgA1 and N327 in IgA2_m1/n_. It was observed that the IgA C-terminal tailpiece shows the highest abundance of sulfated *N-*glycans.

The abundance of sulfated *N-*glycans substantially differs comparing two commercially available human serum IgA samples. In addition to the sulfated *N-*glycan FA2G2S2-SO_4_ (HexNAc_4_Hex_5_Fuc_1_NeuAc_2_Sulfo_1_) described by our previous *N-*glycomic analyses (22, 23), here also other *N-*glycan compositions carrying sulfated HexNAc and *O-*acetylated sialic acid were identified on human IgA. *O*-acetylated *N*-glycans have been also previously characterized in our group by Cajic *et al*. (23), but in horse serum. In the future, the methodology we have developed here will allow site-specific examination of sulfated *N-*glycans in IgA extracted from other body fluids (e.g., saliva or milk) (24). Longitudinal studies that integrate IgA sulfated *N-*glycans can be beneficial in various clinical conditions such as inflammatory bowel diseases or rheumatoid arthritis (17, 36, 37). Expanding the research on the role of sulfated *N-*glycans in IgA effector function can determine whether sulfated *N-*glycans are relevant as a critical quality attribute (CQA) of recombinantly expressed IgA for therapeutic use. Finally, this work also presents an oxonium-ion-guided strategy for finding the parameters necessary to identify the *N-*glycopeptides of interest in IgA. This optimization strategy can be adapted to other instances, and it is particularly useful in cases where the identification of *N-*glycopeptides is hindered by a lack of knowledge about the peptide and *N-*glycan components.

## Experimental Procedures

### Experimental Design and Statistical Rationale

A visualization of the experimental design is presented in the Figure 1. Our work focuses on the reliable site-specific identification of HexNAc-sulfated *N-*glycans on human serum IgA, primarily the FA2G2S2-SO_4_ *N-*glycan, reported via glycomic analysis (22, 23). To this end, the IgA heavy and light chains were separated, and their glycopeptide-enriched fractions were analyzed via MS/MS. An oxonium-ion-guided strategy was used to detect MS^2^ spectra derived from sulfated *N-*glycopeptides. The correct software-assisted identification of the sulfated *N-*glycopeptides was supervised using different search parameters, glycan compositions, and protein modifications. To demonstrate the robustness in the identification of sulfated *N-*glycans, two commercially available human serum IgA samples were analyzed without fractionation. To report on newly discovered *N-*glycan compositions, *de novo* sequencing was conducted on the corresponding *N-*glycopeptide MS^2^ scans primarily acquired at a low MS/MS fragmentation energy. To account the variability caused by the glycopeptide enrichment, the LC-MS/MS measurement, and the different sources of IgA, we conducted cotton-hydrophilic interaction liquid chromatography-solid phase extraction (cotton-HILIC-SPE) and LC-MS/MS measurement for each IgA sample by four replicates. (*N*-glyco)peptide searches were performed implementing defined oxonium ions MS/MS filters and 1% false-discovery rate (FDR).

**Figure 1.**
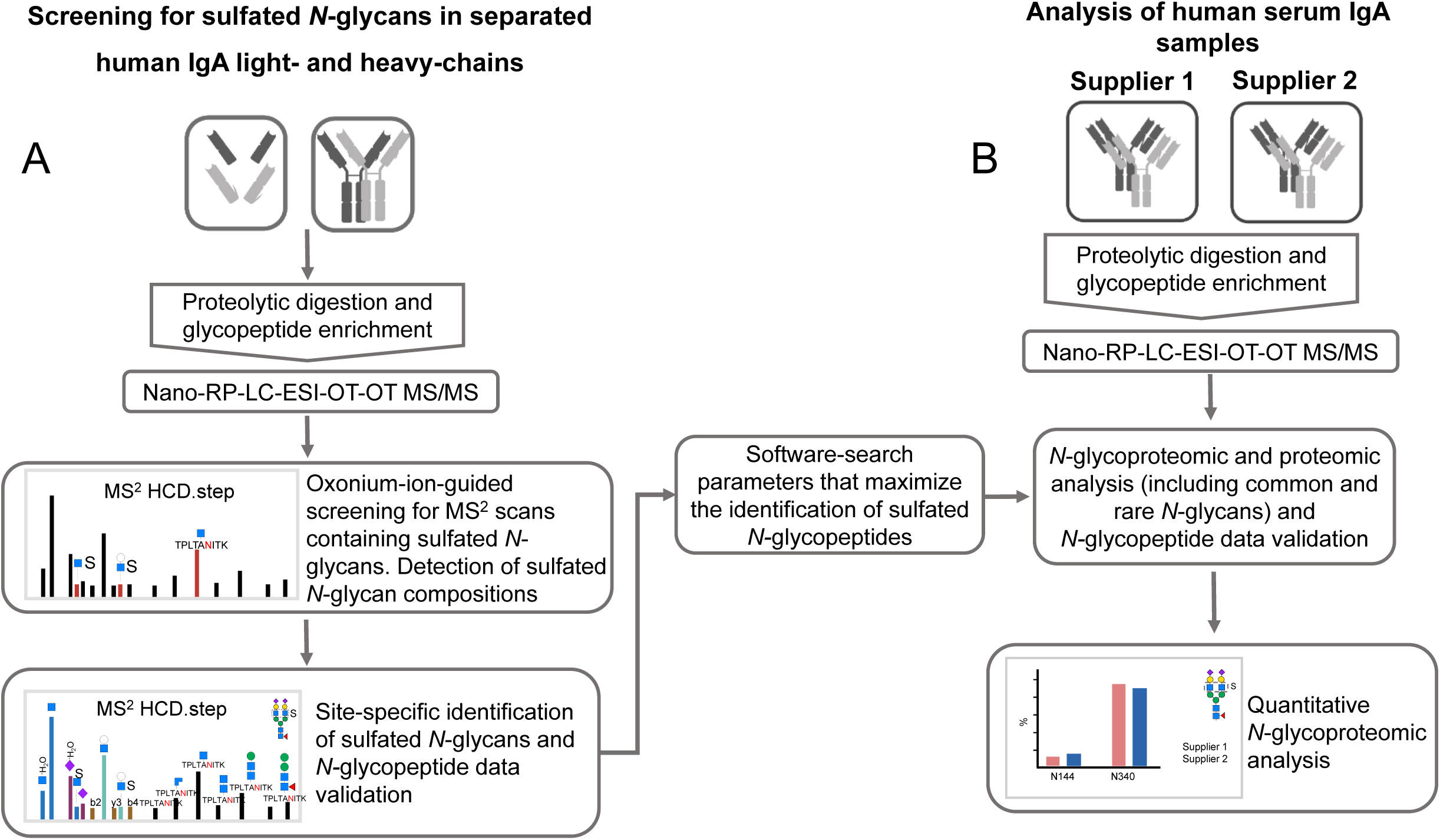
Experimental design for screening for sulfated *N-*glycopeptides. (A) Optimized Workflow. (B) Application of the optimized workflow on two human serum IgA samples. Nano-RP-LC-ESI-OT-OT-M/MS: nano-reversed phase-liquid chromatography with electrospray-ionization coupled orbitrap tandem mass spectrometric measurement for precursor and fragment ions.

The relative abundance of IgA1 and IgA2 subclass in each sample was estimated via a proteomic analysis allowing 1% FDR on peptide and protein level. A label-free quantitative glycoproteomic analysis was conducted on the manually validated *N-*glycopeptides of each IgA sample. The mean and standard deviation of the *N-*glycan relative abundance per *N-*glycosylation site were calculated for the *N-*glycopeptides where at least three technical replicates returned an area under the curve value.

### Samples and Materials

Commercial human IgA purified from blood serum was purchased from two suppliers: supplier 1, Sigma-Aldrich (14036-1MG, St. Louis, MO, USA); and supplier 2, Athens Research & Technology (16-16-090701, Athens, Georgia, USA). Milli-Q water suitable for LC-MS analysis was freshly obtained from Millipore Milli-Q® Advantage A10 system (18,2 MΩ×cm, <5 ppb) equipped with a LC-Pak® Polisher filter unit (#LCPAK0001) purchased from Merck Millipore (Darmstadt, Germany). The reagents applied were MS grade or the highest purity available. LC-MS grade acetonitrile (ACN, A955-212) and trifluoroacetic acid (TFA, 28904) were purchased from Fisher Scientific (Schwerte, Germany). Ammonium bicarbonate (ABC, 09830), formic acid (FA, 56302), DL-dithiothreitol (DTT, D5545), iodoacetamide (IAA, I1149) and calcium chloride (CaCl_2_, A4689) were purchased from Merck (Darmstadt, Germany). Sequencing grade trypsin LC-MS grade was purchased from Promega (#V5111, Madison, WI, USA).

### Protein fractionation by gel electrophoresis

A GELFrEE® 8100 fractionation system, (Abcam, Cambridge, UK) was employed for protein fractionation. The cartridge (8% Cartridge Kit, #42103, Abcam, Cambridge, UK) was set on the system with HEPES 1X running buffer according to the manufacturer instructions. Four chambers were loaded with IgA (25 μL/chamber equivalent to 25 μg of protein) in reducing and denaturing conditions. The electrophoresis was programmed to generate 12 fractions by controlling time and voltage and conducted according to the manufacturer instructions. The fractions were stored at -20 °C. The 12 protein fractions were evaluated via SDS-PAGE 8-16% Tris-Glycine gel. The first fraction showed a protein band of around 25k Da (IgA L-fraction). Fractions 7–9 were pooled since a band between 60 and 70 kDa was consistently observed (IgA H-fraction). The fractions 2–6 and 10–12 were discarded since no protein band was observed.

### Digestion

Only half of the volume of both IgA fractions (L- and H-) was required for the glycoproteomic analysis (the other half was stored for future experiments). Using the filter aided sample preparation approach (FASP) developed by Wisniewski *et al*. and modified by Hoffman *et al.*, each sample volume was loaded on the 10 kDa Nanosep® Omega Filters (OD010C35, Pall®)(34, 38). The proteolytic digestion of the IgA fractions was conducted as described by Hoffmann *et al*. (34). For digestion, the ratio set was 1 μg of trypsin per 60 μg of protein sample. After adding trypsin, the sample fractions were incubated over night at 37 °C and 300 rpm. In addition, 100 μg of IgA samples from supplier 1 and supplier 2 were digested without prior protein fractionation separately using the same enzyme ratio and incubation conditions. To recover the eluates, the filters were centrifuged at 10,000 x g for 15 min (set up applied for all the following steps). The filter membrane was washed with 50 μL of 50 mM ABC_(aq)_ with 5% (v/v) ACN and then with 50 μL of water. The eluates were dried in a rotational vacuum concentrator (0.01 mbar, ca. 3 h, 1 °C, same parameters were applied for all drying steps).

### Cotton-hydrophilic interaction liquid chromatography-solid phase extraction (Cotton-HILIC-SPE)

The glycopeptide enrichment method applied here (Zuniga-Banuelos *et al.* (35)) is a modification of the method from Selman *et al.* (39). The total amount of tryptic digest from the IgA L- and H-fractions was resuspended in 20 μL 85% (v/v) ACN_(aq)_. The tryptic peptides from the two unfractionated IgA samples (from supplier 1 and 2) were resuspended in 100 μL 85% (v/v) ACN_(aq)_. For each unfractionated sample, the glycopeptide enrichment was conducted by four replicates using 20 μL of tryptic peptides each time. Four HILIC-fractions were collected: depletion, wash, elution 1 and elution 2. Depletion and wash fractions were pooled since both contain almost no glycopeptides. All HILIC-fractions were dried, stored at -20 °C, and resuspended in 10 μL water on the day of injection.

### LC-MS/MS analysis

Nano-reversed-phase liquid chromatography coupled to electrospray ionization orbitrap tandem mass spectrometry (nanoRP-LC-ESI-OT-OT-MS/MS) was conducted using a Dionex UltiMate 3000 RSLCnano system (UHPLC, Thermo Fisher Scientific, Germering, Germany) coupled to an Orbitrap Eclipse Tribrid mass spectrometer. A C18 trap column (length 2 cm, pore size 100 Å, particle size 5 µm, inner diameter 100 µm, Acclaim PepMap™ 100, #164199 Thermo Fisher Scientific, Lithuania), together with a C18 separation column (length 25 cm, pore size 100 Å, inner diameter 75 μm, particle size 2 μm, Acclaim PepMap™ RSLC nanoViper #164941, Thermo Fisher Scientific, Lithuania) were used for sample separation in the UHPLC system. For the run, 3 μL of sample were injected isocratically using the loading pump (100% mobile phase A: 0.1% (v/v) FA_(aq)_) and a flow rate of 7 μL/min. Five minutes after injection the trap column was switched in line with the separation column. The nano pump was operated at a flow rate of 300 nL/min at 40 °C using mobile phase A (0.1% (v/v) FA_(aq)_) and mobile phase B (80% (v/v) ACN_(aq)_ + 0.1% (v/v) FA_(aq)_)to obtain the following separation gradient: 5% B (0–5 min); 5–31% (5–35 min); 31–44% (35–40 min); 44–95% (40–41 min); 95% (41–46 min); 95–5% (46–47 min); 5% (47–60 min). The ion sources spray voltage was static set to 2.55 kV and the temperature of the ion transfer tube was set to 275 °C.

Mass spectrometric data acquisition of precursor (MS or MS^1^) and fragment ion spectra (MS/MS or MS^2^) was conducted with an Orbitrap Eclipse Tribrid mass spectrometer (OT-OT-MS/MS; Thermo Fisher Scientific, Dreieich, Germany) at higher-energy collisional dissociation (HCD) with stepped normalized collisional energy (NCE 20, 35, 50 %) “HCD.step”, as described by Hoffmann *et al.* (34). MS precursor ion scans were performed in positive ion mode using a scan range of m/z 350 to 1,500. The automatic gain control (AGC) target was set to “standard” with a resolution of 120,000 and a maximum injection time of 50 ms. Dynamic exclusion was set as follows: exclusion duration to 10 sec, repeat count one, and mass tolerance of 10 ppm. Data-dependent acquisition mode with 1.5 sec cycle time between precursor spectrum acquisitions. MS data-dependent fragment ion scans were acquired performing HCD.step at a resolution of 60,000, scan range set to “normal” and isolation window set to m/z 1.6.

Additional acquisition of fragment ion scans was triggered by the detection of at least two out of the three characteristic oxonium marker ions (HexNAc_1_Hex_1_ [M+H]^+^/ 366.1395, HexNAc_1_Sulfo_1_ [M+H]^+^/ 284.0435, HexNAc_1_Hex_1_Sulfo_1_ [M+H]^+^/ 446.0963) with mass accuracy of 10 ppm. Also here, an HCD method with a fixed NCE of 20 “HCD.low” was employed at a resolution of 60,000 with a scan range set to “normal” and isolation window of m/z 1.6. Resulting from the different modified fragmentation regimes, HCD.low fragment ion scans complemented the information content of the acquired HCD.step spectra (34).

### Oxonium-ion guided detection of MS^2^ spectra containing sulfated HexNAc and identification of new *N*-glycan compositions

The MS^2^ spectra were screened for sulfated HexNAc oxonium marker ions (HexNAc_1_Sulfo_1_ [M+H]^+^/ 284.0435 and Hex_1_HexNAc_1_Sulfo_1_ [M+H]^+^/ 446.0963) using a scan filter in Thermo Xcalibur Qual Browser software (Version 2.2, Thermo Fisher Scientific, Bremen, Germany) as described by Zuniga-Banuelos *et al.* (35). The mass of the sulfated *N-*glycan moieties was estimated for each MS^2^ scan by subtracting the observed peptide mass from the precursor mass. The putative peptide mass was determined by using a conserved fragmentation pattern: (1) [M_peptide_+H -NH_3_]^+^, (2) [M_peptide_+H]^+^, (3) [M_peptide_+H +^0.2^X HexNAc]^+^ and (4) [M_peptide_+H +HexNAc]^+^ (NH_3_=17.0265 Da, ^0.2^X HexNAc=83.0371 Da; and HexNAc=203.0794 Da) (34). The putative peptide and *N*-glycan masses list for the selected MS^2^ spectra is presented in Supplementary Table 1. The compositions of the sulfated *N-*glycans were predicted by entering the estimated mass of the sulfated *N-*glycan moiety in the open source GlycoMod tool from expasy.org (40).

### *N-*Glycoproteomic and proteomic data analysis

The first data analysis (Figure 1A) produced a comprehensive identification of the *N-*glycopeptides bearing sulfated *N-*glycans in the HILIC-fractions derived from the IgA H-fraction (fraction showing evidence for sulfated *N-*glycans). The (*N-*glyco)proteomic data analysis included three steps: 1) identification of *N-*glycopeptides harboring sulfated *N-*glycans, 2) identification of *N-*glycopeptides harboring common *N-*glycans, and 3) identification of non-glycopeptides. Multiple glycoproteomic analyses using Byonic™ search engine (v4.4.2 by Protein Metrics, Cupertino, CA, USA) were done to explain the HCD.step MS^2^ spectra containing sulfated *N-*glycans. Supplementary Table 2 describes the parameters for all searches applied on the three steps. Primarily, four protein sequences were used on the *N-*glycoproteomic searches: IgA1 and IgA2 heavy constant chains, J-chain and pIgR. All the (*N-*glyco)proteomic searches were conducted with Byonic™. The results from the three searches and all raw files were imported in Byologic™ software (v4.6-37-gda29558119 x64, Protein Metrics, Cupertino, CA, USA) to create one (*N-*glyco)peptide identification list, which was then manually validated.

The second data analysis round (Figure 1B) focused on the technical replicates from the two IgA samples (from two suppliers) not fractionated by GELFrEE® system. For each sample, one proteomic and four glycoproteomic searches were conducted using Byonic™. Both proteomic and *N-*glycoproteomic searches included the MS/MS raw data from all HILIC-fractions (depletion/wash, elution 1, and elution 2). The proteomic searches were set using the human canonical proteome UniprotKB (June 2022, 20386 canonical sequences). The parameters set for the proteomic and *N-*glycoproteomic search per supplier sample are shown in Supplementary Table 3. There it is also described that three dedicated *N-*glycoproteomic searches supported the identification of following *N-*glycan modifications: 1) *N-*glycan compositions with HexNAc sulfation, 2) *N-*glycan compositions with *O-*acetylated sialic acid, and 3) *N-*glycan compositions with glucuronic acid. Four additional searches per sample focused on the identification of *N*-glycopeptides from truncated variants of the IgA1- and IgA2-tailpiece. These searches are also described in Supplementary Table 3. The results from the 18 searches were imported and combined in Byologic™, to create one (*N-*glyco)peptide identification list. The *N-*glycopeptide identifications with a “Byonic MS2 search score” above 100 were manually validated (as described in Zuniga-Banuelos *et al.* (35)) and considered for further analyses. The non-glycopeptide identifications with a “Byonic MS2 search score” above 100 were trusted and considered for further analyses without manual validation.

### Relative quantification of (*N-*glyco) proteomic data

The relative abundance of IgA1 and IgA2 was calculated on all technical replicates of the IgA samples from both suppliers by including the MS/MS raw files obtained from all HILIC-fractions. Label-free quantification was conducted within a single proteomic analysis using Proteome Discoverer (version 2.5.0.400, Thermo Fisher Scientific, Bremen, Germany). The software setup allowed defining the files by supplier and technical replicates before the analysis. The search engines Sequest HT (Proteome Discoverer 2.5.0.400, Thermo Fisher Scientific, Bremen, Germany) and Mascot (version 2.6, Matrix Science, London, UK) were set using the human protein database SwissProt/UniprotKB (20,315 canonical sequences, v2022-06-14), full specific tryptic digestion, two missed cleavages allowed, precursor and fragment mass tolerance: 10 ppm and 0.02 Da respectively. The following dynamic modifications were set: deamidated (N, Q), oxidation (M), acetyl (protein N*-*Terminus). Carbamidomethyl (C) was set as static modification. Percolator was applied for peptide validation allowing 1% FDR on peptide level.

Minora feature detector node, included in the processing workflow, enabled detecting and grouping peptide-signals for the HILIC-fractions that belong to the same technical replicate. The label-free quantification applied in the consensus workflow integrated the nodes feature mapped and precursor ions quantifier, considering “unique+razor” peptides with all other parameters set as default.

The proteomic data derived from Proteome Discoverer was processed with Microsoft Excel (2016). The relative abundance of IgA subclasses per replicate was calculated by normalizing the integrated peak area of each representative IgA1- or IgA2-peptide (“NFPPSQDASGDLYTTSSQLTLPATQCLAGK” or “NFPPSQDASGDLYTTSSQLTLPATQCPDGK”, respectively) including variants due to missed cleavages and chemical modifications by the total integrated peak area of both peptides. The values of the technical replicates were averaged for each supplier sample.

Label-free quantification was also carried out on the validated (*N-*glyco)peptide list from the second Byonic™ data analysis. The (*N-*glyco)proteomic data combined in Byologic™ was processed with Microsoft Excel (2016) for relative quantification of *N-*glycan compositions per site (micro-heterogenity). The (*N-*glyco)peptides were classified by *N-*glycosylation site, peptide sequence homology, and then grouped by *N-*glycan compositions (neglecting differences caused by peptide modifications and missed cleavages). The *N-*glycan compositions were quantified if at least three technical replicates showed values from valid *N-*glycopeptide identifications. The abundance of each *N-*glycan composition was normalized by the total area under the curve of all *N-*glycan compositions found per corresponding *N-*glycosylation site. The values of the technical replicate were averaged for each supplier sample.

## Results and Discussion

Site-specific *N-*glycans on the Fc region from each human IgA subclass play a particular role in the effector function. Our recently published glycomic experiments have demonstrated the presence of *N-*glycans bearing sulfated HexNAc in human serum IgA (22, 23). However, the specific position and the relative abundance of sulfated *N-*glycans in human serum IgA subclasses have not been elucidated so far. Here, an in-depth *N-*glycoproteomic workflow, previously established by us (35), was applied to human serum IgA and optimized for identifying the site-specific position of the sulfated *N-*glycan FA2G2S2-SO_4_, previously reported by Cajic *et al.* and Chuzel *et al.* (22, 23). The MS^2^ spectra of each fractionated IgA subunit (heavy and light chain) were screened for HexNAc-sulfated oxonium ions, in order to start the optimization. A rationalized evaluation of selected MS^2^ spectra led to determining the software-assisted search parameters needed for identifying glycopeptides bearing HexNAc-sulfated *N-*glycans. This process maximized the identification of sulfated *N-*glycans on human serum IgA, uncovering not only the position of the previously reported sulfated *N-*glycan but also five additional HexNAc-sulfated *N-*glycan compositions. These compositions were all present in the MS^2^ spectra derived from the IgA heavy chain fraction analysis. To showcase the efficacy of the optimized workflow, the *N-*glycoproteomic analysis was applied to two commercially available human serum IgA samples. Our approach generated a reliable site-specific identification of *N-*glycans modified not only with sulfation, but also with *O-*acetylation. This enabled a comprehensive comparison of micro-heterogeneity of the *N-*glycosylation of these two human serum IgA samples.

### Oxonium-ion guided detection of MS^2^ spectra containing sulfated HexNAc

To maximize the identification of sulfated *N-*glycopeptides on IgA, the light and heavy chains of human serum IgA were reduced, denatured and fractionated (Supplementary Figure 1); both fractions were subsequently subjected to tryptic digestion and cotton-HILIC-SPE glycopeptide enrichment. The obtained glycopeptide enriched fractions were analyzed by mass spectrometry (MS) and fragmented using stepped collisional energy. Both, precursor and fragment ions were measured by high-resolution mass spectrometry. The data analysis firstly focused on identifying sulfated *N-*glycans in the MS^2^ spectra. To this end, the MS^2^ spectra containing sulfated HexNAc oxonium marker ions (“sulfo-glycopeptide spectra”) were screened. These screenings revealed 32 “sulfo-glycopeptide spectra” exclusively in the glycopeptide-enriched IgA fraction (Supplementary Table 1), apparently composed of IgA heavy chain subunits (Supplementary Figure 1). Sulfated *N-*glycopeptides were not found in the IgA fraction expected to contain the J- and light-chains of IgA1 and IgA2. This screening supports the assumption that the observed sulfated *N-*glycans are most likely attached to the heavy chains of the IgA subclasses (26, 27). In addition to the sulfated *N-*glycan HexNAc_4_Hex_5_Fuc_1_NeuAc_2_Sulfo_1,_ (reported as FA2G2S2-SO_4_ in our previous glycomic studies (22, 23)), masses corresponding to other sulfated *N-*glycans were identified and their compositions were proposed using GlycoMod tool, as described in the Experimental Procedures section of this work (40). These new sulfated *N-*glycan compositions were integrated in the following *N-*glycoproteomic search.

### Identification of sulfated *N-*glycopeptides in the IgA heavy chain fraction

One hypothesis for the source of the sulfated *N-*glycan observed in glycomic analyses was that it was originally attached to any of the *N-*glycosylation sites of the heavy chain constant region of serum IgA subclasses. Another hypothesis considers that the sulfated *N-*glycans were rather attached to J-chain or pIgR proteins – these two proteins form structural conformations with IgA in its dimeric form and it is reported that this form is present in low abundance in the blood serum (2). In order to evaluate these hypotheses, the amino acid sequences of the IgA1 and IgA2 heavy chains, J-chain and PIgR were employed in all searches. Our above presented result, the detection of “sulfo-glycopeptide spectra”, allowed us to monitor the application of the different *N-*glycoproteomic search strategies. This approach allowed the determination of the optimal search settings to match all previously detected “sulfo-glycopeptide spectra”. As a result, the optimal search parameter settings are shown in Supplementary Table 2. The Table also shows additional search strategies applied to identify *N-*glycopeptides with common *N-*glycans and non-glycopeptides. The resulting combined *N-*glycopeptide identifications were validated, as described in Zuniga-Banuelos *et al.*, and are displayed in the Supplementary Table 4 (35).

The presence of HexNAc-sulfated oxonium marker ions was confirmed in the (*N*-glyco)peptide spectra matches (gPSMs) containing sulfated *N-*glycans assigning the category “HexNAc-Sulfo ion” (Supplementary Table 4). As summarized in the Supplementary Table 1, 29 “sulfo-glycopeptide spectra” could be assigned to 12 different sulfated *N-*glycopeptides with the tailpiece site (N340-IgA1/N327-IgA2_m1/n_). Two more “sulfo-glycopeptide spectra” were assigned to one sulfated *N-*glycopeptide bearing the C_H_2 domain site N205-IgA2_m1/m2/n_. One “sulfo-glycopeptide spectrum” displayed a poor number of fragment ions, thus it was not possible to identify the corresponding sulfated *N-*glycopeptide. Figure 2 shows b and y ion evidence of a peptide bearing a sulfated *N-*glycan at the tailpiece, compared to an identical glycopeptide without sulfation. We also observed that most gPSMs with the tailpiece site N340-IgA1/N327-IgA2_m1/n_ harbor the sulfated *N-*glycan HexNAc_4_Hex_5_Fuc_1_NeuAc_2_Sulfo_1_, which was structurally confirmed by *de novo* sequencing using HCD.step MS^2^ spectra (Figure 2A). Comparing fragment spectra of both, HexNAc-sulfated and non-sulfated *N-*glycan forms in Figure 2B, the B and Y ions supporting this structural difference (in the terminal HexNAc) are evident.

**Figure 2.**
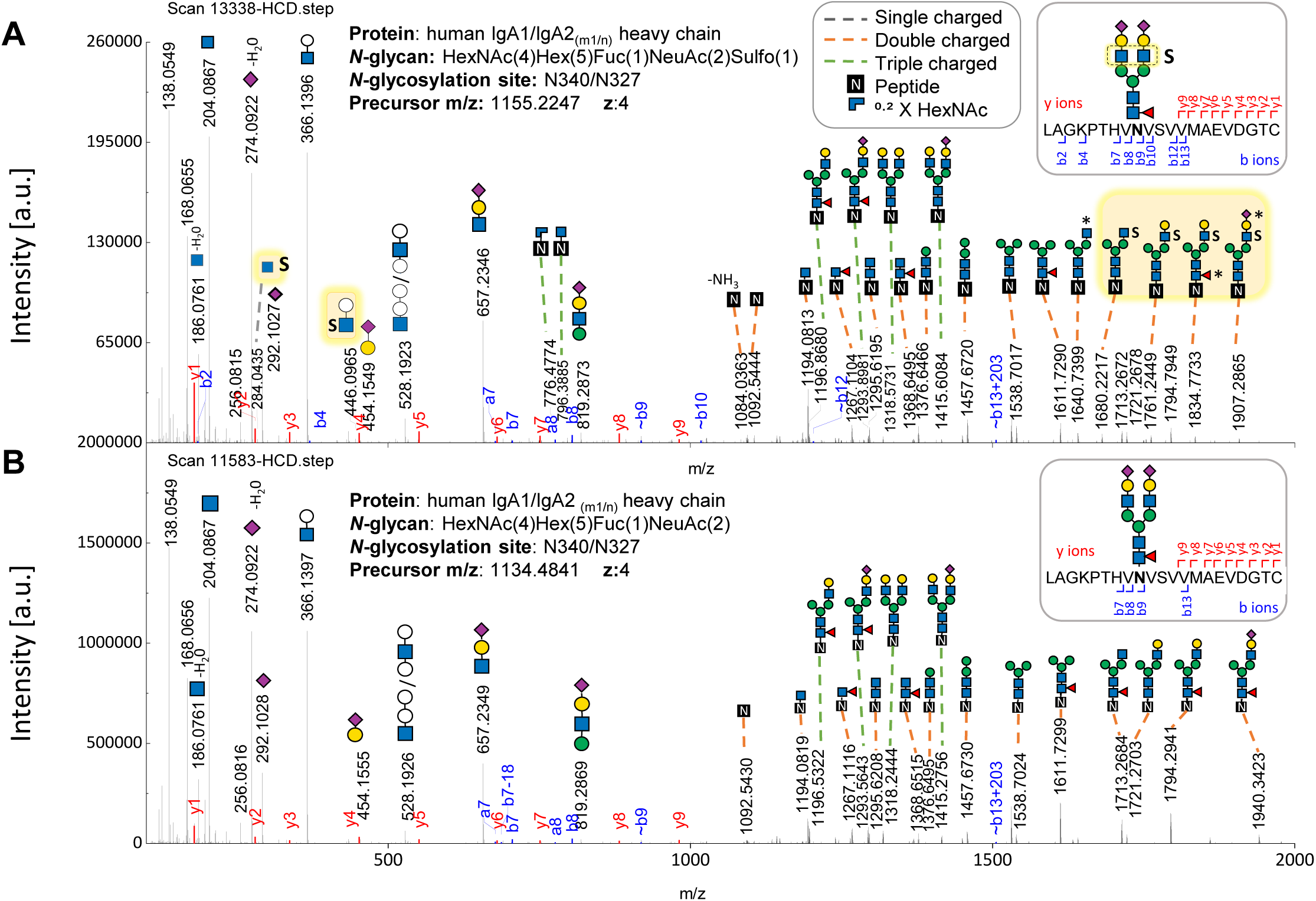
Comparison of HCD.step fragment ion spectra derived from two *N-*glycopeptides (with the same peptide backbone) -one bearing the sulfated versus the one bearing the non-sulfated *N-*glycan at the *N-*glycosylation site in the tailpiece. (A) HCD.step spectrum shows B and Y ions confirming the sulfated *N-*glycan HexNAc(4)Hex(5)Fuc(1)NeuAc(2)Sulfo(1). B and Y ions supporting structural evidence on sulfated *N-*glycans highlighted in yellow. (B) HCD.step spectrum shows the B and Y ions observed in the common non-sulfated *N-*glycan HexNAc(4)Hex(5)Fuc(1)NeuAc(2).

In regard to the J-chain and the pIgR proteins, the (*N-*glyco)proteomic searches did neither show sulfated nor common *N-*glycopeptides from the J-chain, but only peptides (Supplementary Table 4). In addition, the searches resulted in only one gPSM with a non-sulfated complex-type *N-*glycan at the N469 site of pIgR. Overall, this result demonstrates that all sulfated *N-*glycans detected in the IgA fraction analyzed can be assigned to specific *N-*glycosylation sites in the heavy chain constant region of both IgA subclasses, so that the second hypothesis can be rejected. While other glycoproteomic works only have associated galactose-sulfated *N-*glycans to the heavy chain region from IgA subclasses, yet without a site-specific description (25–27), the site-specific identification of sulfated *N-*glycans in IgA is shown here for the first time.

### Peptide modifications detected in the sulfated *N*-glycopeptides identified in the IgA heavy chain fraction

A key adjustment for the optimization of the *N-*glycopeptide search was the number of peptide chemical modifications. Multiple gPSMs with the tailpiece *N-*glycosylation site N340-IgA1/N327-IgA2_m1/n_ correspond to different mass variations of the predicted tryptic peptide (Supplementary Tables 1 and 4). We found that these shifts in peptide mass were caused by one or two amino acid oxidations, one missed cleavage, or a truncated tailpiece form in which the C-terminal tyrosine is absent – this tyrosine truncated tailpiece form has been previously reported (13, 15, 41). Although methionine was the most frequent oxidized amino acid, the software also identified aspartic acid, lysine, and proline oxidation. Oxidation of these amino acids may be caused by oxidative stress in the blood plasma (42–44). Interestingly, one gPSM containing the tailpiece site was identified by a semi-specific tryptic search, thus matching a C-terminal ragged peptide (LAGKPTHVNVS.V) lacking the last 11 amino acids of the tailpiece (VVMAEVDGTCY). Studies have demonstrated that the penultimate cysteine (Cys352) is critical for IgA dimerization, and disulfide bond formation with J-chain and other proteins (45–47). Clarifying whether this short peptide results from an unspecific cleavage or it is another truncated tailpiece variant of the IgA subclasses requires further investigation.

### New HexNAc-sulfated *N-*glycans discovered with the analysis of the IgA heavy chain fraction

Besides the peptide chemical modifications, the generated gPSMs showed six different sulfated *N-*glycan compositions at the tailpiece site N340-IgA1/N327-IgA2_m1/n_. In addition to the previously reported sulfated *N-*glycan, FA2G2S2-SO_4_ (22, 23), five new *N-*glycan compositions, for which sulfated HexNAc oxonium marker ions were detected, could be identified. The structures of the new sulfated *N-*glycan compositions were described to some extent, through their HCD.low and HCD.step fragment ion spectra. The Supplementary Figures 2–6 present the manual annotation of these five new sulfated *N-*glycans. These spectra revealed one HexNAc-sulfated di-fucosylated hybrid-type *N-*glycan, two HexNAc-sulfated sialylated complex-type *N-*glycans without core fucosylation, and one HexNAc-sulfated, sialylated, bisected *N-*glycan with core fucosylation. A fifth sialylated and core fucosylated complex-type *N-*glycan holding two sulfated monosacharides was identified; while sulfated HexNAc is evident, it was not possible to confirm the position of the second sulfation (Supplementary Figure 4).

In follow-up, the GlyConnect *N-*glycan databases corresponding to IgA1 and IgA2_m2_ (UniProt entries: P01876 and P01877, respectively) were screened for sulfated *N-*glycan compositions (48, 49). Interestingly, both Glyconnect *N-*glycan databases showed only two of the six sulfated *N-*glycan compositions detected by us: HexNAc_4_Hex_5_Fuc_1_NeuAc_2_Sulfo_1_ and HexNAc_4_Hex_5_NeuAc_2_Sulfo_1_ (Supplementary Table 5) (25–27). However, these two *N-*glycans were reported to be modified with galactose sulfation instead of GlcNAc sulfation. As reported by She *et al.*, the detection of galactose sulfation in intact *N-*glycopeptides relies on the manual identification of Y ions holding terminal galactose sulfation (50). Our data analysis does not provide any probative structural information on galactose sulfation, for several reasons. First, this type of identification requires establishing a reliable and less time-consuming computational strategy for the identification of the expected Y ions (50). Second, the oxonium marker ion Hex_1_Sulfo_1_ [M+H]^+^, potentially generated by a sulfated galactose, was never detected in any of our gPSMs containing sulfated *N-*glycans. Third, even though detecting only the oxonium marker ion Hex_1_HexNAc_1_Sulfo_1_ [M+H]^+^ implies either a Hex or a HexNAc sulfation, this was the case for only two gPSM and they were verified. The verification revealed that these gPSM were part of the same chromatographic peak as other gPSM presenting HexNAc_1_Sulfo_1_ [M+H]^+^ ion, indicating that these two gPSMs most likely hold sulfated HexNAc as well.

In a minority of cases, the manual validation of the *N-*glycoproteomic results revealed that sulfated *N-*glycan compositions were assigned to some gPSMs without showing sulfated HexNAc oxonium marker ions. However, these gPSMs demonstrated a correct match in regard to the peptide fragment ions (b and y ions) and common glycan oxonium ions (e.g. HexNAc_1_ [M+H]^+^, NeuAc_1_ [M+H]^+^, HexNAc_1_Hex_1_ [M+H]^+^). It is likely that these gPSMs correspond to very low abundant precursor ions or to other sulfated *N-*glycans, including sulfated galactose.

Although manual validation can pursue the identification of galactose-sulfated *N-*glycans, software tools capable of comprehensively Y ion matching can facilitate this identification process. For example, GlycanFinder software performs *de novo* sequencing of *N-*glycopeptides and can be customized to identify peptides bearing galactose-sulfated *N-*glycans (51).

### Application of the optimized *N*-glycoproteomic analysis to two human serum IgA samples

To evaluate the application of the optimized *N-*glycoproteomic search, human serum IgA samples from two commercial suppliers were directly analyzed (not fractionated by GELFrEE® system) and compared. To this end, the tryptic digested samples from both suppliers were enriched for glycopeptides. Technical quadruplicates were obtained for each commercial sample by repeating the cotton-HILIC-SPE protocol four times. All HILIC-fractions from the technical replicates (depletion/wash, elution 1, and elution 2) were analyzed by nanoRP-LC-ESI-OT-OT-MS/MS. All measurements of the HILIC-fractions were included in the proteomic and *N-*glycoproteomic analyses.

As described in Experimental Procedures, we calculated the relative abundance of each IgA subclass based on the quantification of selected peptides identified by Proteome Discoverer. The peptide identification list is provided in Supplementary Table 6. Around 80% of the protein sequence of IgA1 and IgA2 is homologous. Therefore some peptides are assigned to both IgA subclasses, which can lead to misleading results. In order to avoid bias caused by tryptic peptides with homologous IgA sequences (ambiguous peptides), the non-glycopeptides “NFPPSQDASGDLYTTSSQLTLPATQCLAGK” and “NFPPSQDASGDLYTTSSQLTLPATQCPDGK” were selected to represent IgA1 and IgA2, respectively. We found that the ratio between IgA1:IgA2 subclasses was 89:11 in the commercial sample from supplier 1, which is the natural ratio in blood serum (1), and approximately 99:1 in the one from supplier 2, which is ten times higher (Supplementary Table 7). This abnormal ratio in the sample from supplier 2 indicates for a massive enrichment of IgA1, probably due to the purification process applied (52).

In addition, multiple *N-*glycoproteomic searches featuring modified and common *N-*glycan compositions plus a proteomic search using Byonic ™ search engine were conducted (Supplementary Table 3). In order to include (*N-*glyco)peptides from “contaminant” proteins, the human canonical proteome UniProtKB was set as protein database. As described in Experimental Procedures, (*N-*glyco)peptide identifications with a “Byonic MS2 search score” below 100 were excluded due to their poor quality and the remaining *N-*glycopeptide identifications were manually validated. As a result, the Supplementary Table 8 show peptides and *N-*glycopeptides belonging to IgA subclasses and “contaminant” proteins found among all replicates from both samples. The majority of “contaminant” *N-*glycopeptides belongs to α1-antitrypsin, complement C3, kinninogen-1 and α-2-HS-glycoprotein. The presence of these “contaminant” proteins in the purified IgA is probably due to chromatographic co-elution or to stable protein-protein interactions during the purification process (52–54). Studies show that proteins like complement C3 and β-2-glycoprotein form circulating immune complexes with IgA subclasses in blood serum (53, 54). Additionally, a strong interaction is controlled by the penultimate cysteine of the IgA tailpiece forming a disulfide bond either with albumin or α1-antitrypsin (47). Therefore, it is important to consider the contribution of *N-*glycans from “contaminant” proteins in glycomic analysis. For example, oxonium marker ion signals from glucuronidated *N-*glycans were found in the serum IgA samples from both suppliers. However, the *N-*glycoproteomic search revealed that these *N-*glycans do not belong to any *N-*glycosylation site of IgA1 or IgA2 but to α1-antitrypsin and α-2-HS-glycoprotein. Another example is the competitive abundance of sulfated *N-*glycopeptides derived from other proteins. The Supplementary Table 9 shows that, in the case of the sample from supplier 2, the sulfated *N-*glycan HexNAc_4_Hex_5_Fuc_1_NeuAc_2_Sulfo_1_ was also found on *N*-glycopeptides from α-2-HS-glycoprotein with an average relative abundance of 41.4%, normalized to the total amount of this sulfated *N-*glycan present in the sample from supplier 2.

### Description of the site-specific *N*-glycosylation of human serum IgA including sulfated and *O*-acetylated *N*-glycans

Figure 3 displays a comparative relative quantification of the *N-*glycosylation micro-heterogeneity of the human serum IgA samples from both suppliers. Granting that ambiguous peptides represent the IgA tailpiece site (N340-IgA1/N327-IgA2_m1/n_) and the C_H_2 domain site (N144-IgA1/N131-IgA2_m1/m2/n_), the micro-heterogeneity analysis on those *N-*glycosylation sites is assigned to both IgA subclasses (Supplementary Table 10 and 11). The micro-heterogeneity analysis for specific *N-*glycosylation sites of IgA2 is listed separately in Supplementary Tables 12 and 13. As the bar plot displays in Fig. 3A, while the tailpiece site N340-IgA1/N327-IgA2_m1/n_ harbors core fucosylated sialylated complex-type di- and multi-antennary *N-*glycans, the C_H_2 domain site N144-IgA1/N131-IgA2_m1/m2/n_ bears mostly non-fucosylated sialylated hybrid- and di-antennary complex-type *N-*glycans. Thus, the *N-*glycans observed by us, at the sites in the C_H_2 domain and in the tailpiece, were also observed in other glycoproteomic analyses that included the simultaneous analysis of both IgA subclasses, demonstrating the consistency of our micro-heterogeneity analysis with other works (14–17). By separation of IgA1 and IgA2 subclasses, Steffen *et al.* have described that the total IgA2 *N-*glycome shows low levels of sialylated bisected *N-*glycans and higher levels of hybrid- and oligomannose-type *N-*glycans (12). Chandler *et al.* also showed by the specific glycoproteomic analysis of IgA2, that the C_H_2 domain site N131-IgA2_m1/m2/n_ bears hybrid-type *N-*glycans in higher abundance than the C_H_2 domain site N144 in IgA1 (14). These studies suggest that the share of hybrid-type *N-*glycans will be influenced by the IgA1:IgA2 ratio, which might explain the differences observed in our analysis, between both commercial samples. The *N-*glycosylation sites that belong only to the IgA2 subclass (N92-IgA2_m2/n_, N205-IgA2_m1/m2/n_, and N327-IgA2_m2_) are also described in Fig. 3B. It was observed that these sites predominantly show sialylated core-fucosylated complex-type *N-*glycans, including bisecting *N-*glycans. Chandler *et al.* also observed that these *N-*glycans were dominant at the N92-IgA2_m2/n_ and the N205-IgA2_m1/m2/n_ sites (14).

**Figure 3.**
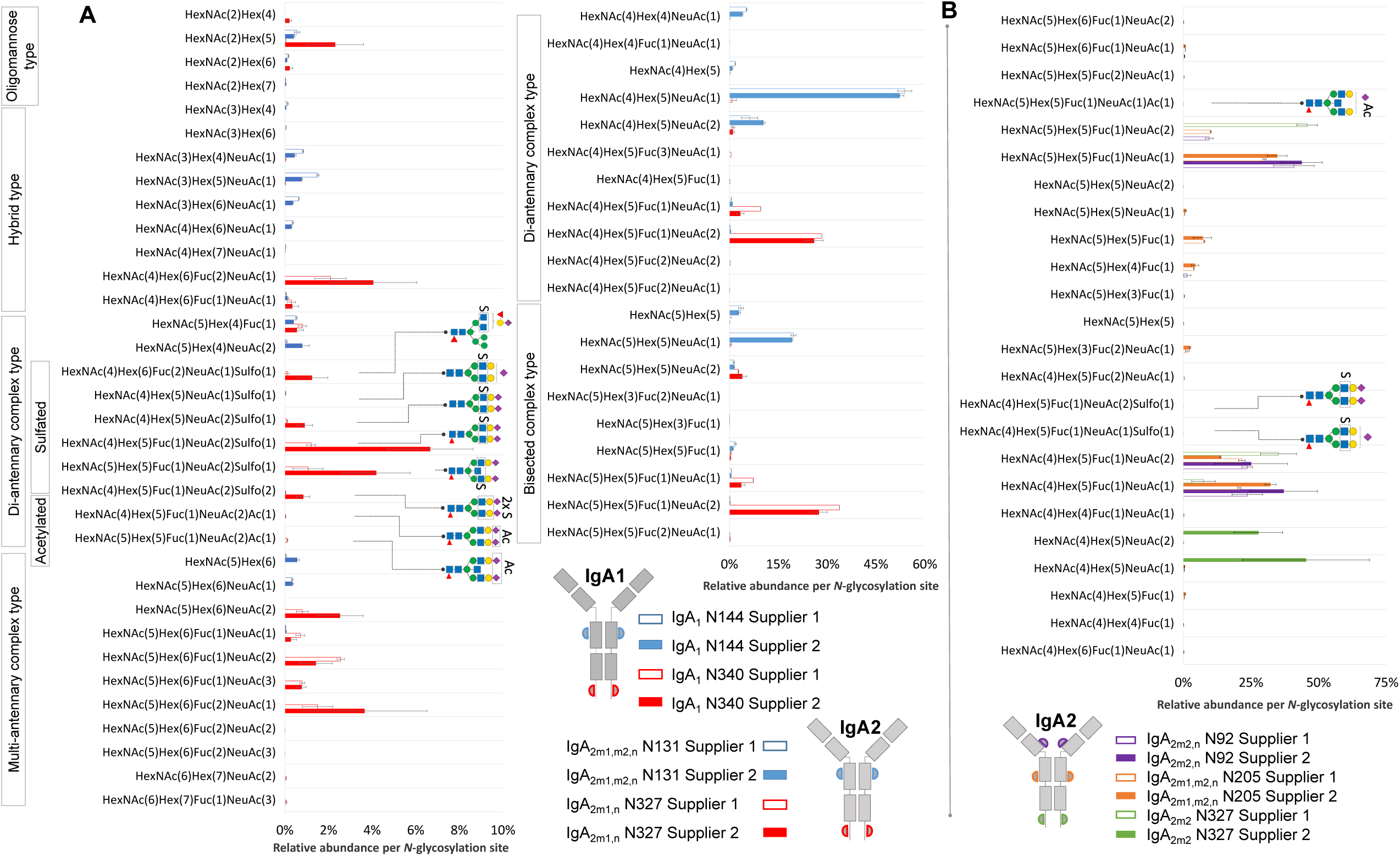
Comparison of micro-heterogeneity of *N-*glycosylation identified in commercially available human serum IgA samples from two suppliers. A) Relative abundance of *N-*glycans at the homologous *N-*glycosylation sites N340-IgA1/N327-IgA2_(m1/n)_ and N144-IgA1 / N131-IgA2_(m1/m2/n)_. B) Relative abundance of *N-*glycans at the *N-*glycosylation sites that are exclusively present in IgA2 subclasses. Results for relative quantification are shown in the Supplementary Tables 10–13.

As Figure 3 shows, the sulfated *N-*glycan HexNAc_4_Hex_5_Fuc_1_NeuAc_2_Sulfo_1_ (previously reported by us via glycomic analyses as FA2G2S2-SO_4_ (22, 23)) was identified with in highest abundance at the tailpiece *N-*glycosylation site N340-IgA1/N327-IgA2_m1/n_ and in a very low abundance at the C_H_2 domain site N205-IgA2 (only 0.04% of the total *N-*glycans in the sample from supplier 1, Supplementary Table 12). At the C_H_2 domain site N144-IgA1/N131-IgA2_m1/m2/n_, the samples from both suppliers bear non-fucosylated sialylated sulfated di-antennary complex-type *N-*glycans in very low relative abundance (Supplementary Table 10 and 11).

The relative abundance of sulfated *N-*glycans varies between the samples from both suppliers (Supplementary Tables 10–13). For example, the bisected complex-type *N-*glycan HexNAc_5_Hex_5_Fuc_1_NeuAc_2_Sulfo_1_ shows 1.06% and 4.19% in the sample from supplier 1 and supplier 2, respectively. In addition, the *N-*glycan HexNAc_4_Hex_5_Fuc_1_NeuAc_2_Sulfo_1_ (FA2G2S2-SO_4_ (22, 23)) displays an average relative abundance of 1.2% and 6.66% in the sample from supplier 1 and supplier 2, respectively. Further analyses are required to elucidate differences in the dominance of sulfated *N-*glycans between the IgA subclasses. Nonetheless, we hypothesize that this substantial difference might be related to the IgA1: IgA2 ratio observed in both IgA samples, where the sample from supplier 2 showed a ten times higher abundance of IgA1. Interestingly, this would suggest that sulfated *N-*glycans are selectively more conserved at the tailpiece *N-*glycosylation site in IgA1 and not in IgA2.

Figure 4 depicts a site-specific description of the sulfated and *O-*acetylated *N-*glycans identified hereby in the human serum IgA samples from both suppliers. As listed in the Supplementary Table 5, one *N-*glycan bearing *O-*acetylation of sialic acid (HexNAc_4_Hex_5_Fuc_1_NeuAc_2_Ac_1_) has been associated to IgA1 (25–27). Thus, a specific *N-*glycoproteomic search including *N-*glycan compositions with *O-*acetylated NeuAc was conducted as described in Experimental Procedures and Supplementary Table 3. This resulted in the identification of such reported NeuAc *O-*acetylated *N-*glycan at the tailpiece site N340-IgA1/N327-IgA2_m1/n_ (depicted in Supplementary Figure 7), plus three new NeuAc *O-*acetylated *N-*glycan compositions.

**Figure 4.**
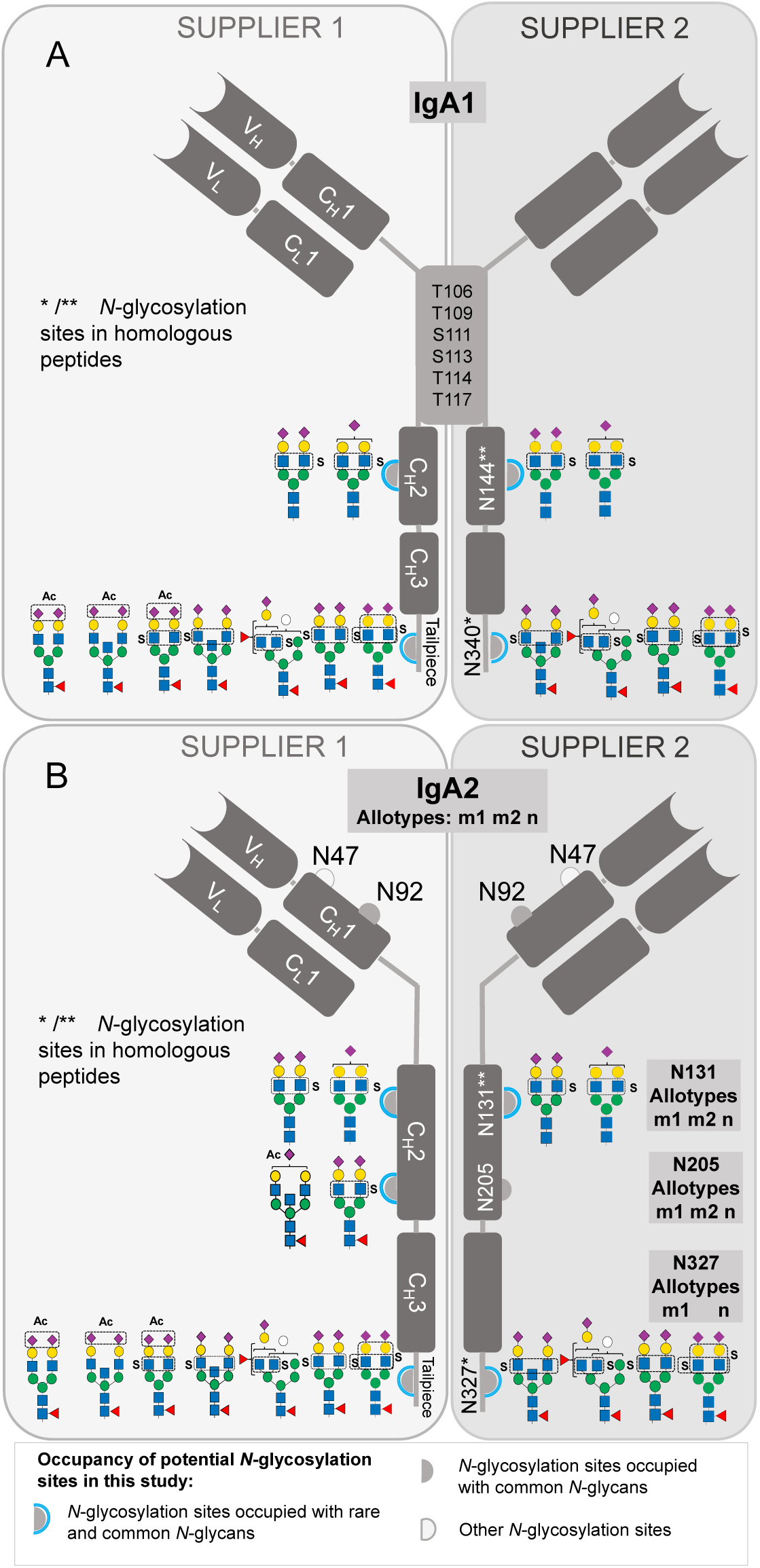
Site-specific description of the here identified *N*-glycans carrying sulfated HexNAc and NeuAc *O*-acetylation in human serum IgA. (A) Site-specific *N-*glycan identification in IgA1 (B) Site-specific *N-*glycan identification in IgA2.

As displayed in Figure 4, all the NeuAc *O-*acetylated *N-*glycans identified here are complex-type, with and without bisecting GlcNAc, core-fucosylated and sialylated. Interestingly, NeuAc *O-*acetylated *N-*glycans were only identified in the human serum IgA sample from supplier 1. The three additional NeuAc *O-*acetylated *N-*glycans detected hereby were *de novo* sequenced to describe them in detail (Supplementary Figures 7–10). One of the NeuAc *O-*acetylated complex-type *N-*glycans was observed at the C_H_2 domain site N205-IgA2_m1/m2/n_, in a very low relative abundance (on average 0.05%, Supplementary Table 12), while the second new NeuAc *O-*acetylated bisected complex-type *N-*glycan was found at the tailpiece site N340-IgA1/N327-IgA2_m1/n_. Together with the non-bisected complex-type *N-*glycan, this second new NeuAc *O-*acetylated bisected complex-type *N-*glycan displays a total relative abundance of 0.14% (Supplementary Table 10). The third additionally identified NeuAc *O-*acetylated complex-type *N-*glycan bears both, NeuAc *O-*acetylated and sulfated HexNAc, and was found at the tailpiece site. Since this third *N-*glycopeptide appeared only in one technical replicate of the sample from supplier 1, it was not part of the quantification. This very rare *N-*glycan was further interpreted by manual *de novo* sequencing in the HCD.low MS^2^ spectrum (Supplementary Figure 10A). This MS^2^ spectrum not only presents oxonium ions from *O-*acetylated sialic acid, but also Hex_1_HexNAc_1_Sulfo_1_ [M+H]^+^ oxonium ion and Y ions demonstrating sulfation at the terminal HexNAc. Moreover, the HexNAc_1_Sulfo_1_ [M+H]^+^ oxonium marker ion was evident in the HCD.step MS^2^ spectrum (Supplementary Figure 10B).

The role of NeuAc *O-*acetylated *N-*glycans in IgA is unknown. Studies focused on *O-*acetylated sialic acid have reported that NeuAc *O-*acetylation may have immunoregulatory effects by affecting the affinity of the sialylated glycan for lectins (e.g. CD22) or bacterial sialidases (55, 56). In addition, NeuAc *O*-acetylation influences the binding to viral neuraminidases during infection (57). However, it requires further efforts to understand the impact of *O*-acetylated *N*-glycan on IgA effector functions.

Our investigation includes for the first time, HexNAc-sulfated and NeuAc *O-*acetylated *N*-glycans in the micro-heterogeneity description of IgA *N-*glycosylation. It was observed that the relative abundance of NeuAc *O-*acetylated *N-*glycans at the tailpiece *N-*glycosylation site is at least 10 times lower than the abundance of sulfated *N-*glycans in the IgA tailpiece within the sample from supplier 1. Overall, both samples consistently showed that the primary source of HexNAc-sulfated *N-*glycans in IgA is the tailpiece *N-*glycosylation site. It is not known how sulfated *N-*glycans influence the effector function of IgA subclasses. On the one hand, sulfated glycoepitopes, are ligands for L-selectin and this interaction plays an important role in cell adhesion and trafficking of immune system cells (31, 32, 58, 59). On the other hand, studies using glycan arrays demonstrate the affinity of specific influenza viruses (and also bacteria) for sialylated glycoepitopes containing sulfated GlcNAc (28, 60, 61). It has also been reported that sialylated *N-*glycans at the IgA tailpiece play a role in the neutralization of influenza virus infection (6). Further, it is possible that IgA sulfated *N-*glycans play also a key role in the IgA antiviral activity for instance by mimicking a cell receptor-ligand used during infection. Also, other IgA effector functions are strongly modulated by the *N-*glycans at the tailpiece. First, *N-*glycosylation positively influences IgA1 binding to complement C3 – complement coated-IgA carries out the complement-coated antibody-transfer (CCAT) mechanism (19, 62). Second, absence of *N-*glycans at the tailpiece site induces the formation of IgA dimers (19). Third, studies have shown that the *N-*glycosylation at the C_H_2 and C_H_3 domains (Fc region) of IgA1 does not affect binding to the FcαRI receptor (13, 18). However, Steffen *et al*. demonstrated that sialylated *N-*glycans on the Fc region can modulate IgA1 effector functions (12).

New methods have been developed for the detailed analysis of sulfated or *O-*acetylated *N-*glycans. Recently, Cajic, S. *et al* established and applied a workflow for multiple analyses of special *N-*glycans by means of removable fluorescent labeling (Fmoc) (23). Using this method, they isolated and exhaustively characterized *O*-acetylated *N*-glycans from horse serum proteins. Another application of this method, was the isolation of a sulfated *N-*glycan from Fmoc-labeled *N-*glycans released from human serum IgA, for further analysis via both MALDI-TOF and xCGE-LIF. The xCGE-LIF glycomic analysis was combined with a highly specific sulfatase and a sulfate-dependent hexosaminidase, both characterized by Chuzel L. *et al* (22). Also, Chuzel *et al*. found that this sulfatase can act as a lectin highly selective for GlcNAc-6-SO_4_ in absence of calcium. Even though each methodology has limitations, cutting-edge glycoanalytical tools can benefit biological and clinical research. Our glycoproteomic workflow can be applied to other proteins to expand the overview of human glycobiology diversity.

## Conclusion

Currently, the difficulty to detect sulfated *N-*glycans by MS-based glycoproteomic approaches, limits the site-specific elucidation of sulfated *N-*glycans attached to proteins, e.g. to IgA subclasses. IgA has clinical relevance due to its multiple roles in the immune system. In this work, we applied our previously developed in-depth *N-*glycoproteomic workflow (35) to study sulfated *N-*glycans on two commercially available human IgA samples isolated from blood serum. With this we achieved the identification of the *N-*glycosylation sites harboring the sulfated *N-*glycan HexNAc_4_Hex_5_Fuc_1_NeuAc_2_Sulfo_1_ (FA2G2S2-SO4), reported up to now only by site unspecific *N-*glycomic investigations (22, 23). We detected new hybrid-type, and complex-type HexNAc-sulfated *N-*glycans in IgA, which were overlooked in previous glycomic and glycoproteomic analyses (12–17, 22, 23). However, we could not detect previously reported galactose-sulfated *N-*glycans in IgA (25–27). Also, we identified new *N-*glycan compositions holding *O-*acetylated NeuAc in a site-specific manner. Finally, we estimated the relative abundance of these sulfated and *O-*acetylated *N-*glycans per glycosylation site and IgA sample. Our data demonstrate for the first time, that the primary protein position for sulfated and *O-*acetylated *N-*glycans is the tailpiece of IgA. The *N-*glycans linked to the tailpiece position of IgA play a substantial role in the defense against infection by sialic-acid-binding virus (6), for the binding to complement C3, and in IgA dimerization (19). Our MS-based *N-*glycoproteomic workflow allows the investigation of very low abundant *N*-glycopeptide forms, like HexNAc-sulfated and NeuAc *O-*acetylated *N-*glycans. This workflow can be applied to other biological samples, e.g. glycoproteins isolated from different body fluids, such as urine, milk, saliva or mucosal secretions. More investigation is required to clarify the function of sulfated and *O-*acetylated *N-*glycans in the protein-glycan interaction and to evaluate if these are relevant e.g. for IgA production as a therapeutic or in the clinical area as a diagnostic tool.

## Acknowledgments

The authors gratefully thank to Valerian Grote for enriching discussions and to Barbara Koehler for her technical support.

## Data availability

The data produced by the proteomic and *N*-glycoproteomic analyses here conducted is available within the supplemental data of this article. The MS raw files were deposited to the ProteomeXChange Consortium, identifier PXD051396, through MassIVE (dataset identifier MSV000094524).

## Supplemental data

This article contains supplemental data.

## Funding

This study was supported by European Commission (EC) Horizon 2020 research and innovation program for F.J.Z.B. and E.R. under the project IMforFUTURE (H2020-MSCA-ITN/721815), and by the Deutsche Forschungsgemeinschaft (DFG, German Research Foundation) for M.H. and E.R. under the project “The concert of dolichol-based glycosylation: from molecules to disease models” (grant identifier FOR2509).

## Author contributions

F.J.Z.B. and E.R Conceptualization; F.J.Z.B methodology; G.L. and F.J.Z.B validation; F.J.Z.B. and M.H formal analysis; F.J.Z.B project administration; F.J.Z.B. and G.L investigation; F.J.Z.B. and G.L. data curation; G.L. and F.J.Z.B visualization; F.J.Z.B writing—original draft preparation; M.H., G.L., U.R. and E.R writing—review and editing; M.H., U.R., and E.R. supervision; U.R. and E.R funding acquisition. All authors have read and agreed to the published version of the manuscript.

## Conflicts of interest

E.R. is founder and CEO of glyXera GmbH. F.J.Z.B is employee of glyXera GmbH and Max Planck Institute. glyXera provides high-throughput glycomic analysis and holds several patents for xCGE-LIF based glycoanalysis. U.R. is shareholder of glyXera GmbH. M.H. and G.L. declare no conflict of interest.

## Abbreviations

IgA: Human immunoglobulin A
S: Sulfation
Ac: *O*-Acetylation
NeuAc: *N*-acetylneuraminic acid
HexNAc: *N*-acetylhexosamine
Hex: Hexose
gPSM: glycopeptide spectra match
HCD.step: higher-energy collisional dissociation with stepped normalized collisional energy [NCE of 20, 35, and 50]
HDC.low: higher-energy collisional dissociation with fixed NCE of 20

