## Supplemental Figures for "Immunoglobulin A Carries Sulfated *N-*glycans Primarily at the Tailpiece Site – An Oxonium-Ion-Guided Approach for Site-Specific *N*-glycan Identification"

### List of Supplementary Figures

**Supplementary Figure 1**. Separation of light and heavy chains of human serum IgA subclasses using the GELFrEE system.

**Supplementary Figure 2.** Description of fragment ions released from a glycopeptide bearing the sulfated *N*-glycan HexNAc(4)Hex(5)NeuAc(1)Sulfo(1).

**Supplementary Figure 3.** Description of fragment ions released from a glycopeptide bearing the sulfated *N*-glycan HexNAc(4)Hex(5)NeuAc(2)Sulfo(1).

**Supplementary Figure 4**. Description of fragment ions released from a glycopeptide bearing the sulfated *N*-glycan HexNAc(4)Hex(5)Fuc(1)NeuAc(2)Sulfo(2).

**Supplementary Figure 5**. Description of fragment ions released from a glycopeptide bearing the sulfated *N*-glycan HexNAc(5)Hex(5)Fuc(1)NeuAc(2)Sulfo(1).

**Supplementary Figure 6**. Description of fragment ions released from a glycopeptide bearing the sulfated *N*-glycan HexNAc(4)Hex(6)Fuc(2)NeuAc(1)Sulfo(1).

**Supplementary Figure 7**. Glycopeptide containing NeuAc *O*-acetylation in the *N*-glycan HexNAc(4)Hex(5)Fuc(1)NeuAc(2)Ac(1).

**Supplementary Figure 8.** Glycopeptide containing the NeuAc *O*-acetylated *N*-glycan HexNAc(5)Hex(5)Fuc(1)NeuAc(1)Ac(1).

**Supplementary Figure 9.** Glycopeptide containing NeuAc *O*-acetylation in the *N*-glycan HexNAc(5)Hex(5)Fuc(1)NeuAc(2)Ac(1).

**Supplementary Figure 10**. Glycopeptide containing NeuAc *O*-acetylation plus HexNAc sulfation in the *N*-glycan HexNAc(4)Hex(5)Fuc(1)NeuAc(2)Ac(1)Sulfo(1).

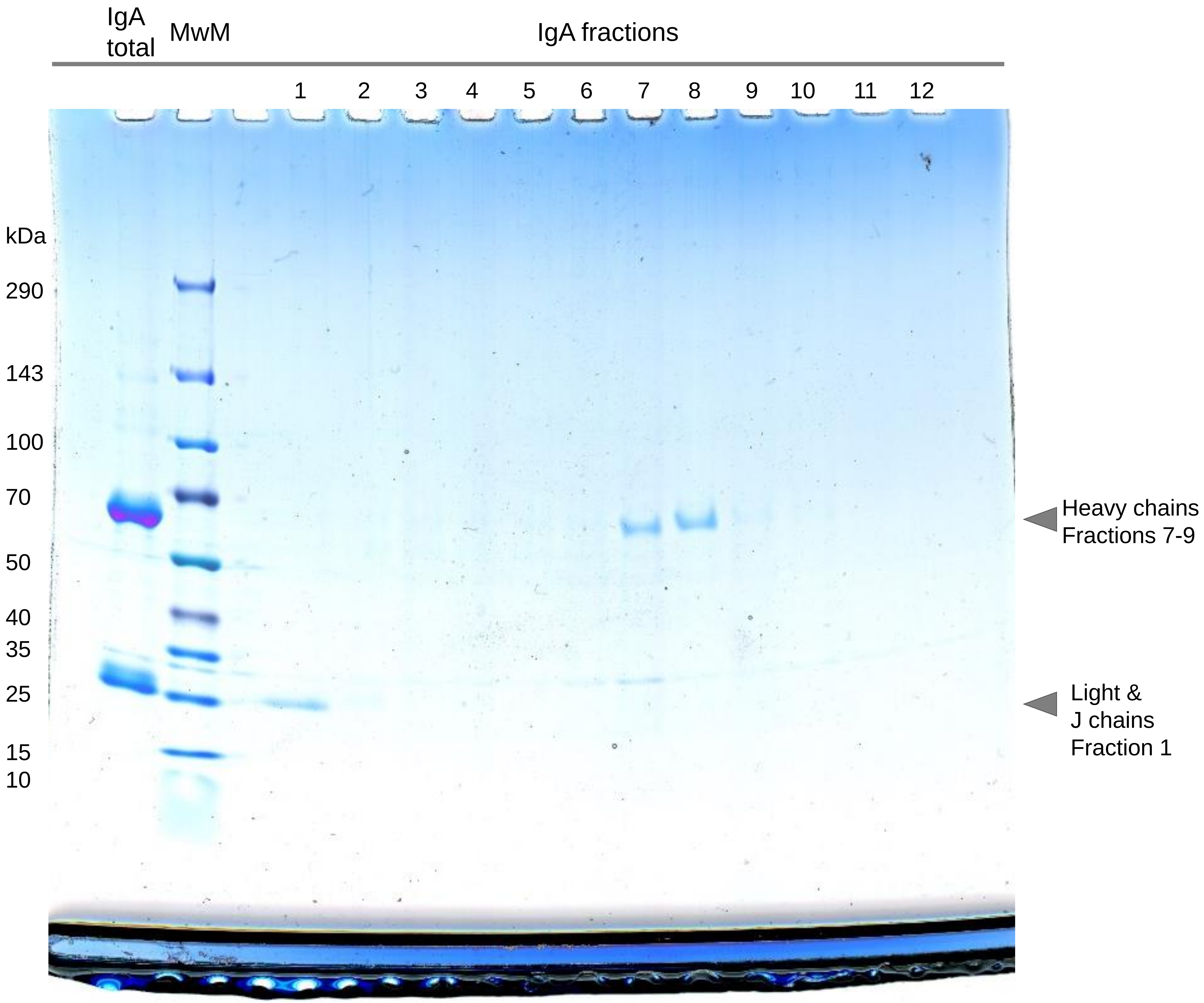


**Supplementary Figure 1. Separation of light and heavy chains of human serum IgA subclasses using the GELFrEE system.** The protein fractionation was evaluated via SDS-PAGE by loading an aliquot of each IgA GELFrEE fraction. MwM: Molecular weight marker.

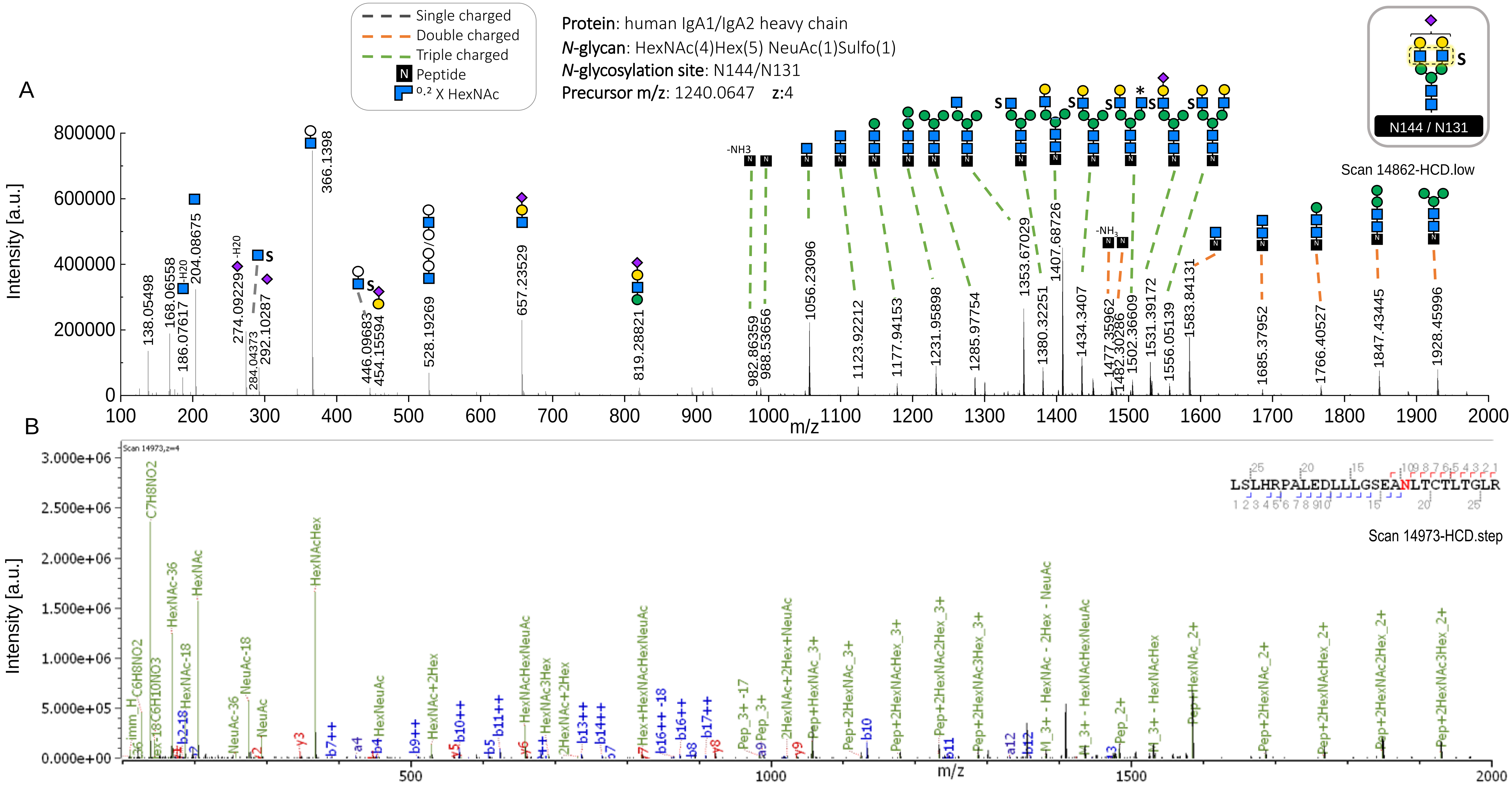


**Supplementary Figure 2. Description of fragment ions released from a glycopeptide bearing the sulfated *N*-glycan HexNAc(4)Hex(5)NeuAc(1)Sulfo(1).** (A) Description of the *N*-glycan moiety through HCD.low spectrum shows B and Y ions confirming antenna with a sulfated HexNAc. The *N*-glycan building blocks with an asterisk symbol (*) depict the *N*-glycan composition by a Y ion assigned to the second isotope peak of the expected m/z. (B) The Byonic ™ annotation of the HCD.step spectrum refers to an amino acid sequence of this *N*-glycopeptide containing the site N144-IgA1/N131-IgA2$_{(m1/m2/n)}$. The HCD.step spectrum shows the oxonium marker ions confirming HexNAc sulfation ($Sulfo_1HexNAc_1$/284.0437 $[M+H]^+$).

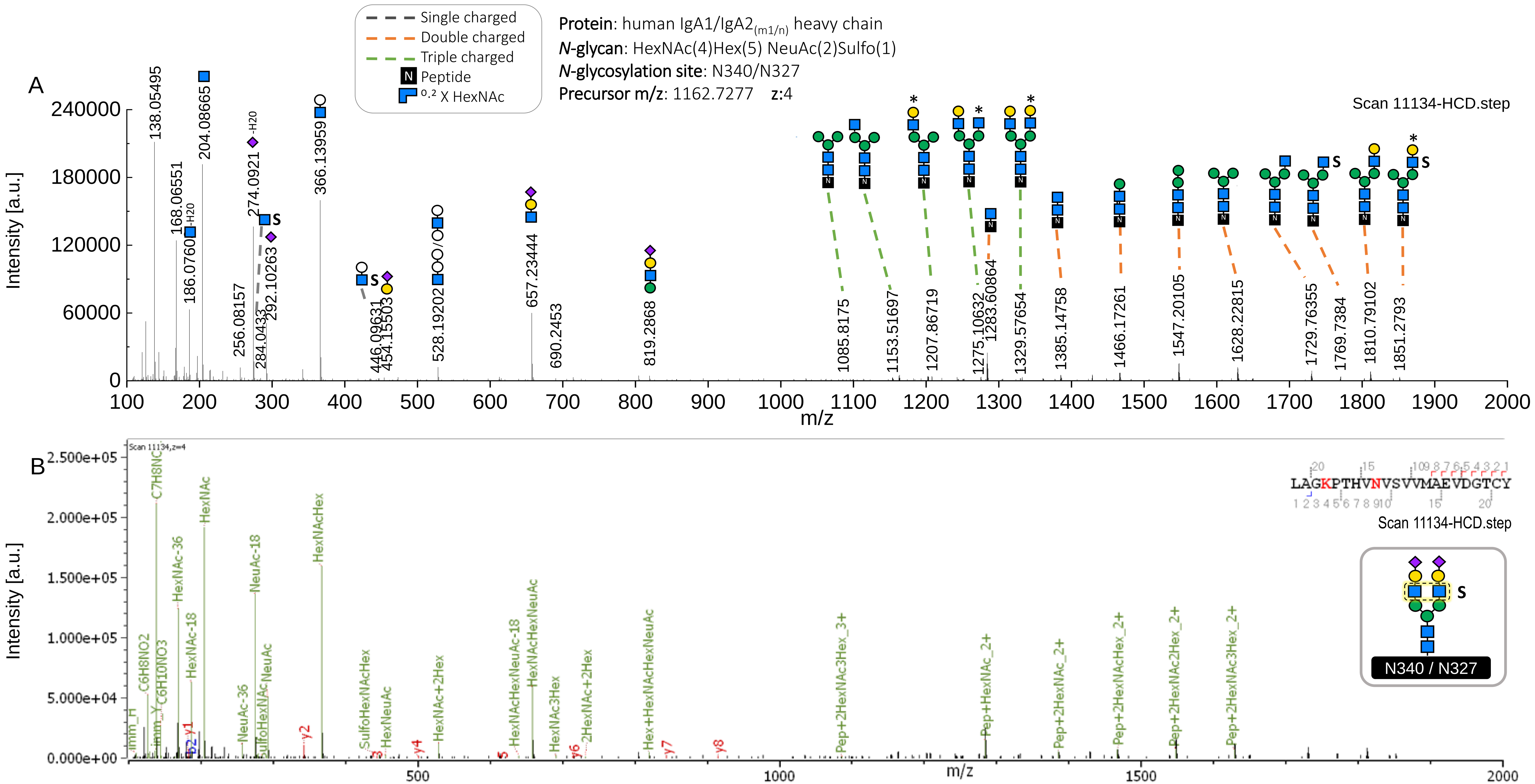


**Supplementary Figure 3. Description of fragment ions released from a glycopeptide bearing the sulfated *N*-glycan HexNAc(4)Hex(5)NeuAc(2)Sulfo(1).** (A) Description of the *N*-glycan moiety through HCD.step spectrum shows B and Y ions confirming antenna with sulfated HexNAc. The *N*-glycan building blocks with an asterisk symbol (*) depict the *N*-glycan composition by a Y ion assigned to the second isotope peak of the expected m/z. (B) Byonic ™ annotation of the HCD.step spectrum refers to an amino acid sequence of this *N*-glycopeptide contains the site N340-IgA1/N327-IgA2$_{(m1/n)}$. The HCD.step spectrum shows the oxonium marker ions confirming HexNAc sulfation (Sulfo$_1$HexNAc$_1$/284.0433 $[M+H]^+$).

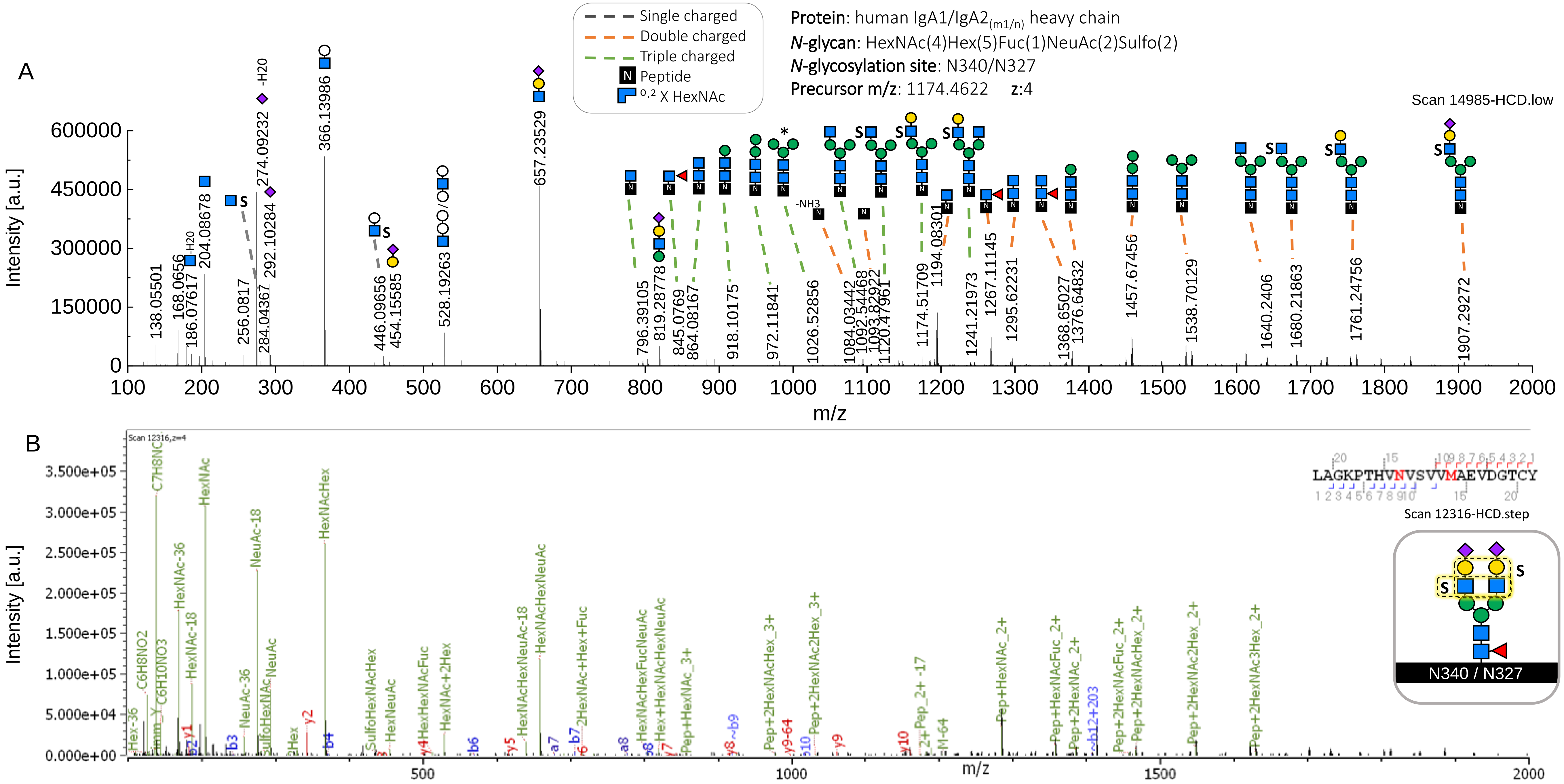


**Supplementary Figure 4. Description of fragment ions released from a glycopeptide bearing the sulfated *N*-glycan HexNAc(4)Hex(5)Fuc(1)NeuAc(2)Sulfo(2).** (A) Description of the *N*-glycan structure through HCD.low spectrum shows B and Y ions confirming one sulfation linked to antenna HexNAc, while the linkage of the second sulfation cannot be fully deduced and it can be associated to antenna HexNAc or Hex. The *N*-glycan building blocks with an asterisk symbol (*) continue to depict the *N*-glycan composition by a Y ion assigned to the second isotope peak of the expected m/z. (B) HCD.step spectrum annotated by Byonic ™ recognizes b and y ions corresponding to an amino acid sequence of this *N*-glycopeptide containing the site N340-IgA1/N327-IgA2$_{(m1/n)}$. The HCD.step spectrum shows Sulfo$_1$HexNAc$_1$/284.0434 $[M+H]^+$ oxonium ion, confirming HexNAc sulfation.

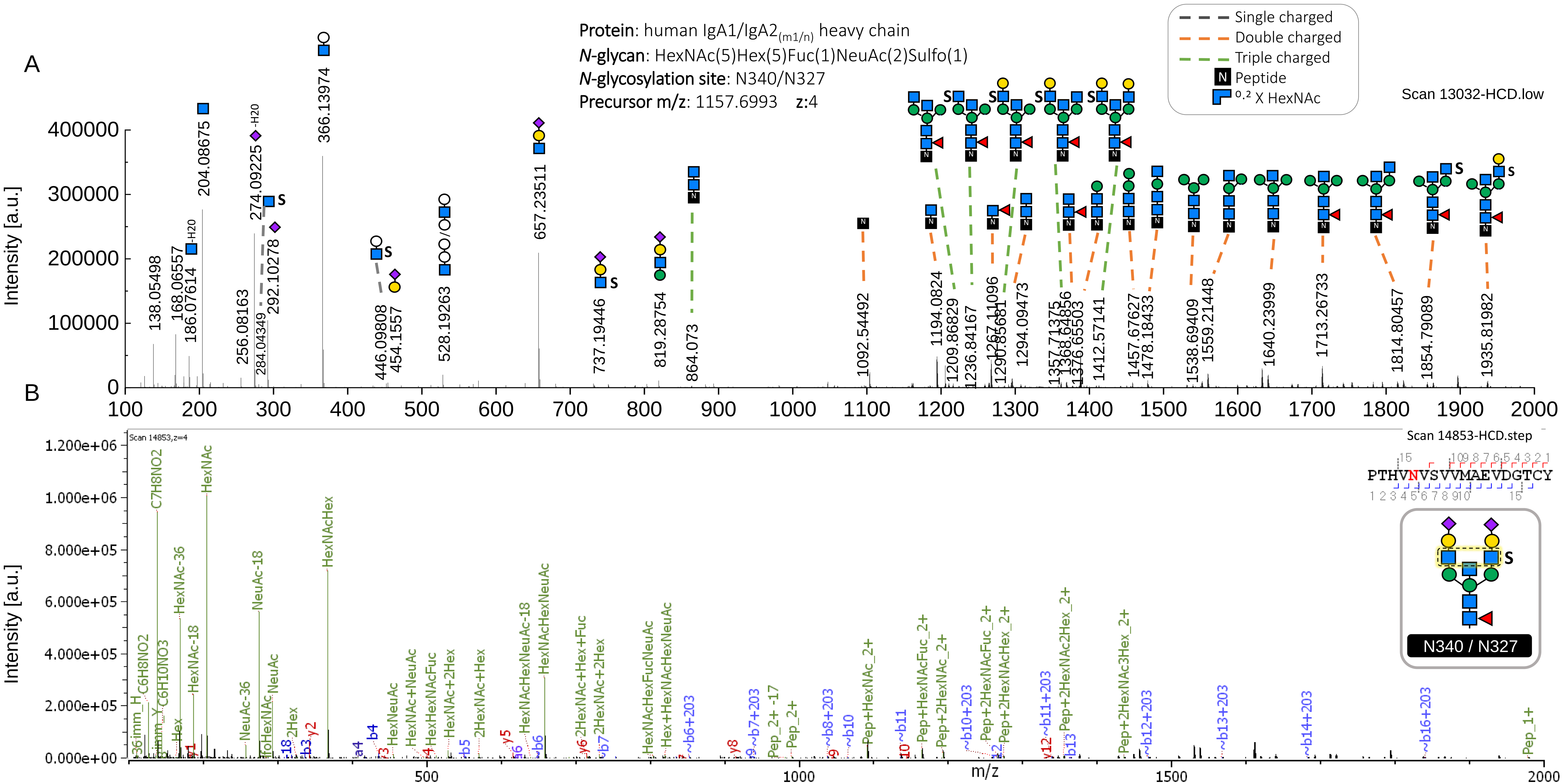


**Supplementary Figure 5. Description of fragment ions released from a glycopeptide bearing the sulfated *N*-glycan HexNAc(5)Hex(5)Fuc(1)NeuAc(2)Sulfo(1).** (A) Description of bisected *N*-glycan structure through HCD.low spectrum shows Y ions indicating sulfation most likely in one antenna HexNAc and not in the bisecting HexNAc. (B) The Byonic ™ annotation of the HCD.step spectrum refers to an amino acid sequence of this *N*-glycopeptide contains the site N340-IgA1/N327-IgA2$_{(m1/n)}$. The HCD.step spectrum also shows the oxonium ion confirming HexNAc sulfation (Sulfo$_1$HexNAc$_1$/284.0439 $[M+H]^+$).

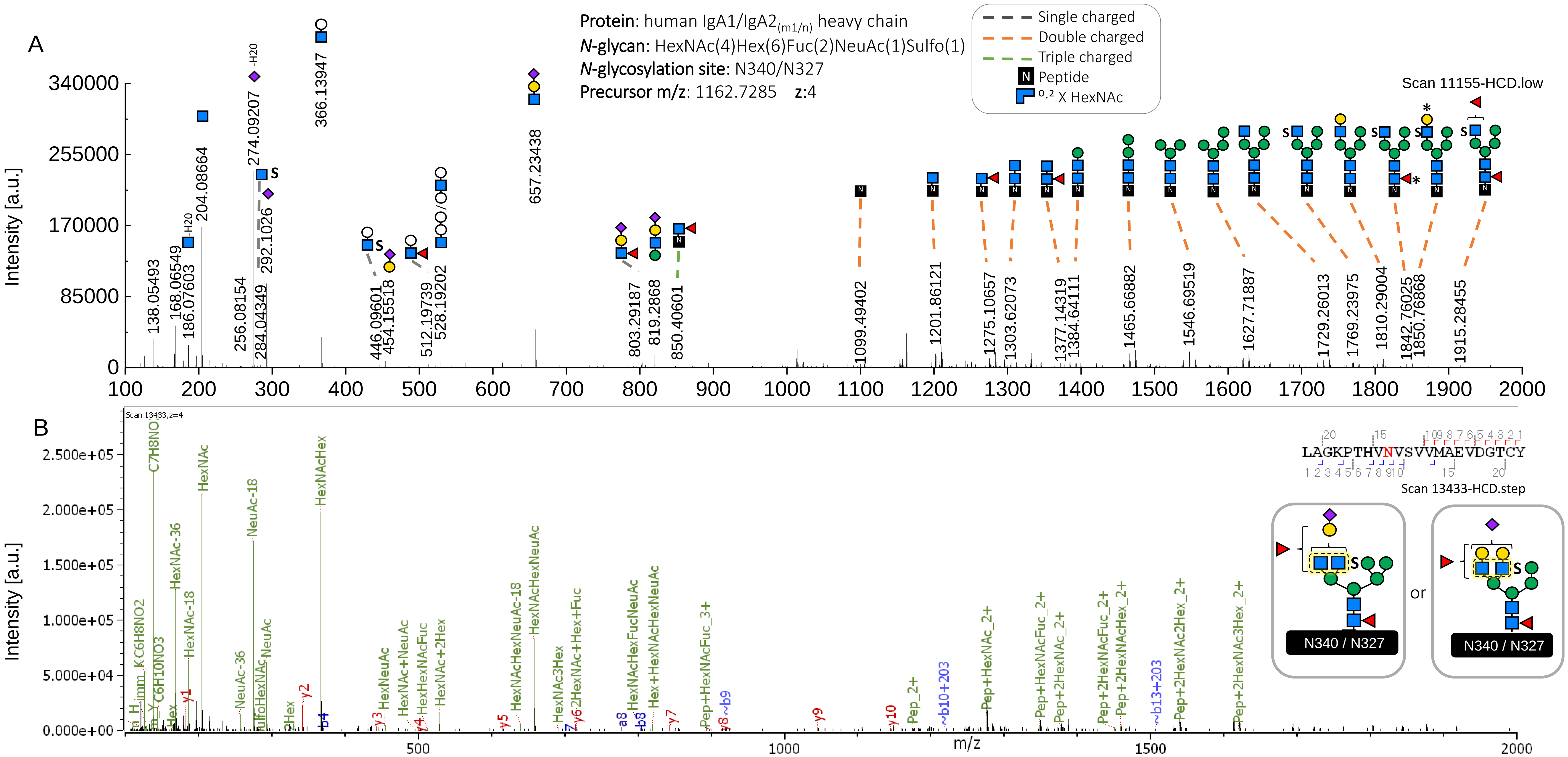


**Supplementary Figure 6. Description of fragment ions released from a glycopeptide bearing the sulfated *N*-glycan HexNAc(4)Hex(6)Fuc(2)NeuAc(1)Sulfo(1).** (A) Description of di-fucosylated hybrid *N*-glycan structure through HCD.low spectrum shows B and Y ions indicating HexNAc sulfation. The *N*-glycan building blocks with an asterisk symbol (*) depict the *N*-glycan composition by a Y ion assigned to the second isotope peak of the expected m/z. (B) Byonic ™ annotation of the HCD.step spectrum refers to an amino acid sequence containing site N340-IgA1/N327-IgA2$_{(m1/n)}$ through the b and y fragment ions released from the *N*-glycopeptide. The HCD.step spectrum shows the oxonium marker ions confirming HexNAc sulfation ($Sulfo_1HexNAc_1$/284.0433 $[M+H]^+$).

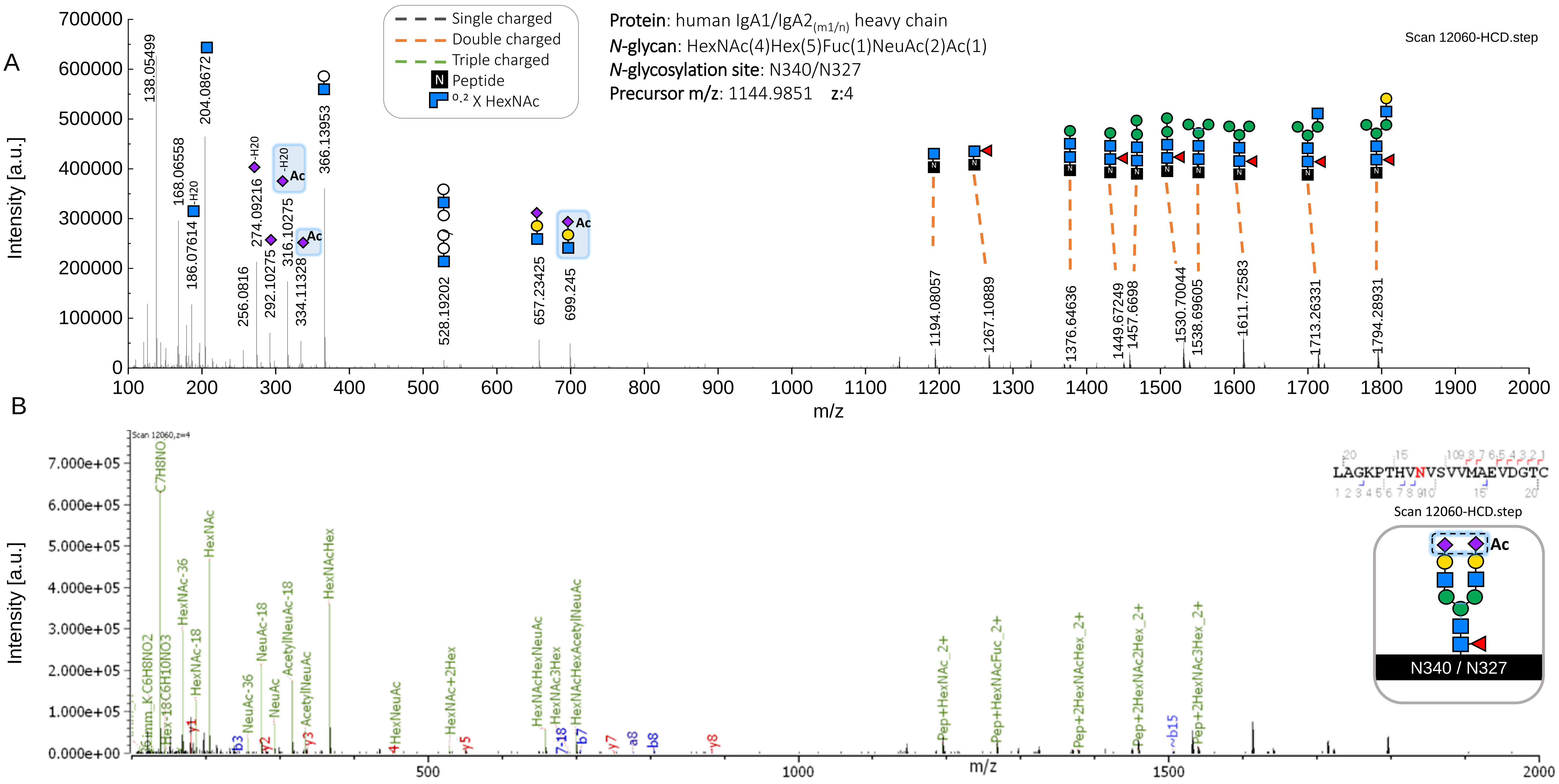


**Supplementary Figure 7. Glycopeptide containing NeuAc *O*-acetylation in the *N*-glycan HexNAc(4)Hex(5)Fuc(1)NeuAc(2)Ac(1).** (A) Description of *N*-glycan structure through HCD.step spectrum shows B and Y ions providing structural evidence on NeuAc *O*-acetylation (highlighted in blue). (B) Byonic ™ annotation of the HCD.step spectrum identifies fragment ions released from a peptide containing the site N340-IgA1/N327-IgA2$_{(m1/n)}$. Oxonium marker ions confirming NeuAc *O*-acetylation (Acetyl$_1$NeuAc$_1$/316.1027 [M-H2O+H]$^+$, Acetyl$_1$NeuAc$_1$/334.1133 [M+H]$^+$ and HexNAc$_1$Hex$_1$Acetyl$_1$NeuAc$_1$/699.2454 [M+H]$^+$).

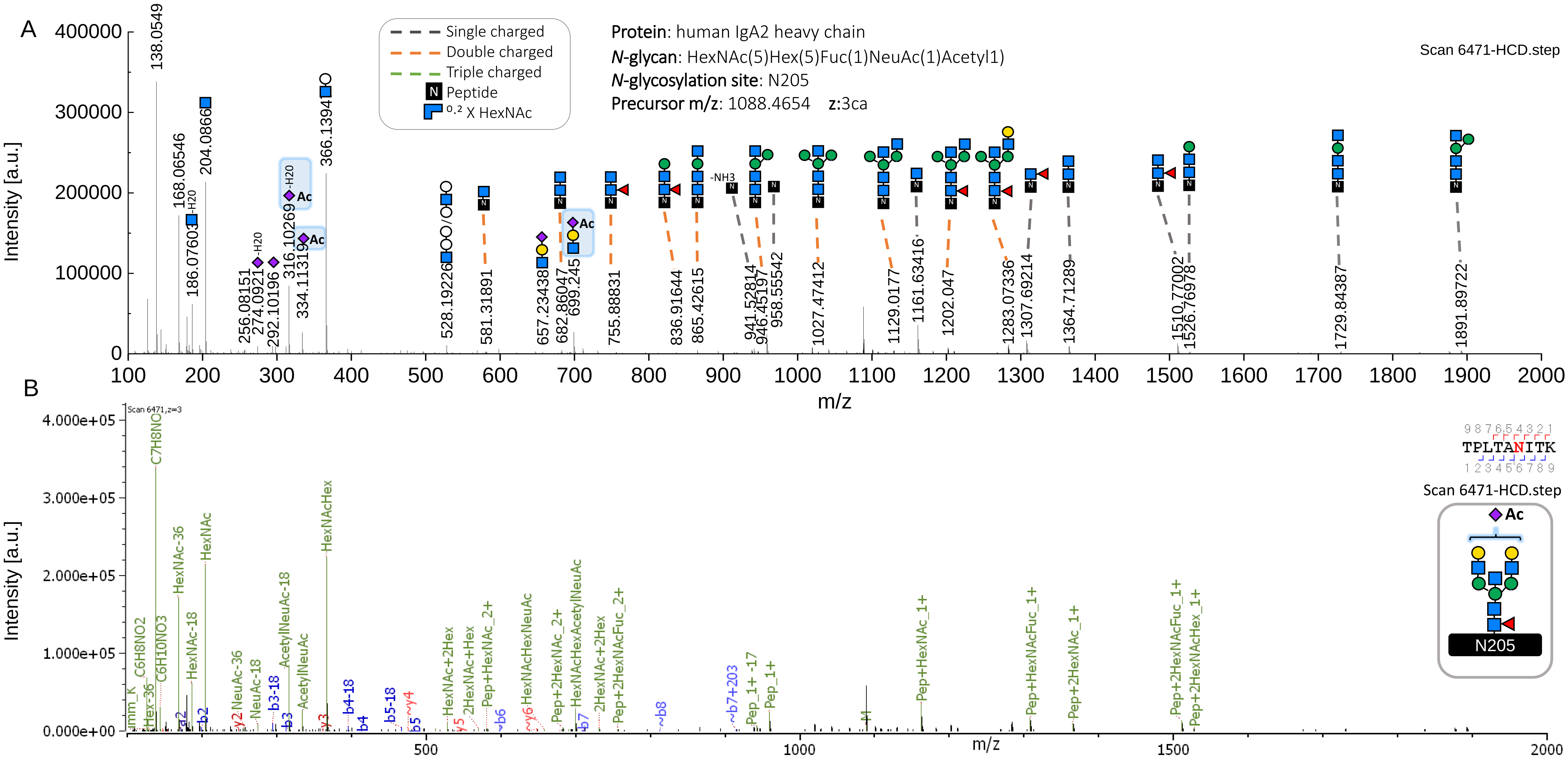


**Supplementary Figure 8. Glycopeptide containing the NeuAc *O*-acetylated *N*-glycan HexNAc(5)Hex(5)Fuc(1)NeuAc(1)Ac(1).** (A) Description of bisected *N*-glycan structure through HCD.step spectrum shows B ions indicating NeuAc *O*-acetylation (highlighted in blue). (B) Byonic ™ annotation of the HCD.step spectrum displays b and y ions that identify an *N*-glycopeptide containing the site N205-IgA2$_{(m1/m2/n)}$. Oxonium marker ions confirming NeuAc *O*-acetylation (Acetyl$_1$NeuAc$_1$/316.1027 [M+H-H$_2$O]$^+$, Acetyl$_1$NeuAc$_1$/334.1132 [M+H]$^+$ and HexNAc$_1$Hex$_1$Acetyl$_1$NeuAc$_1$/699.2450 [M+H]$^+$).

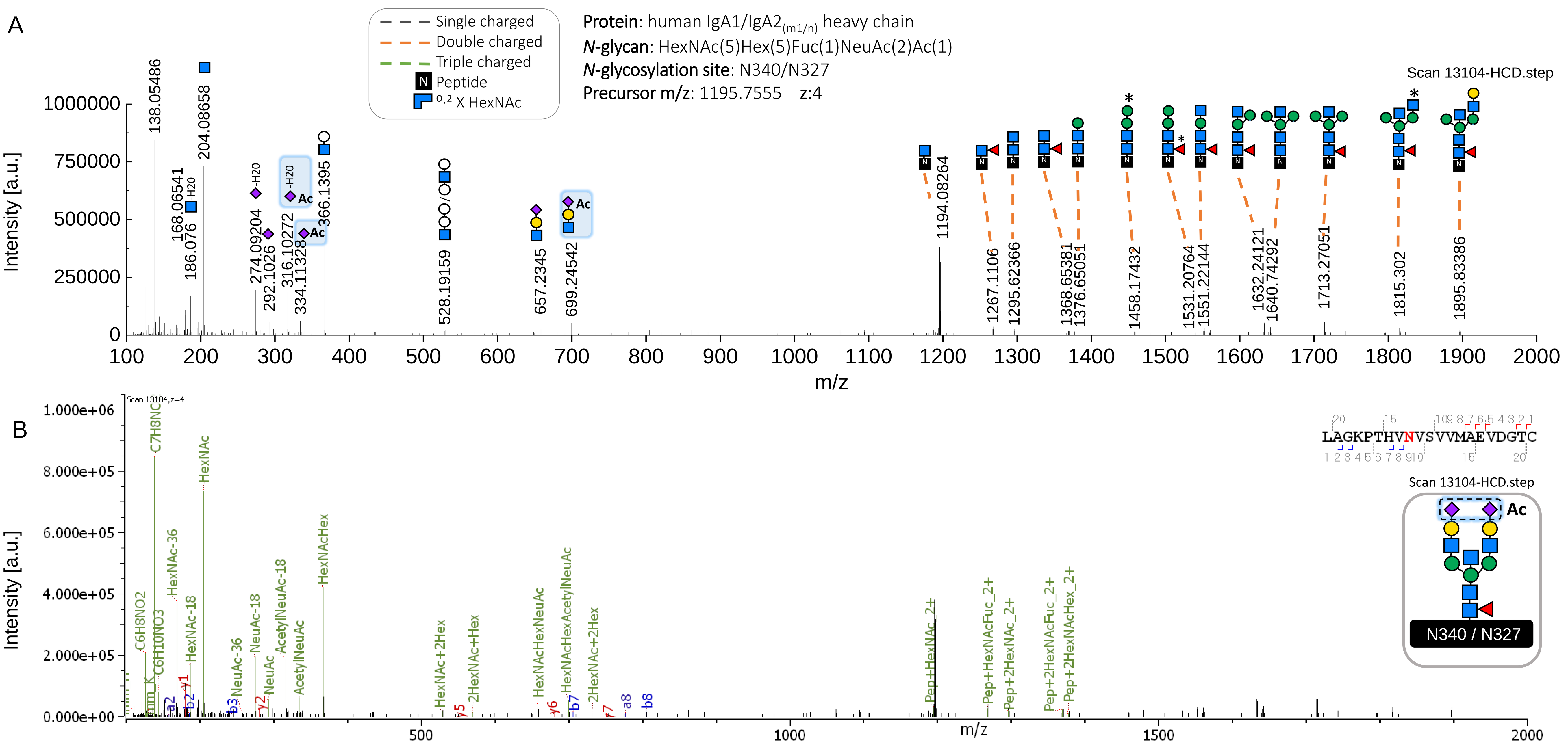


**Supplementary Figure 9. Glycopeptide containing NeuAc *O*-acetylation in the *N*-glycan HexNAc(5)Hex(5)Fuc(1)NeuAc(2)Ac(1).** (A) Description of bisected *N*-glycan structure through HCD.step spectrum shows B ions indicating NeuAc *O*-acetylation (highlighted in blue). The *N*-glycan building blocks with an asterisk symbol (*) depict the *N*-glycan composition by a Y ion assigned to the second isotope peak of the expected m/z. (B) Byonic ™ annotation of the HCD.step spectrum identifies fragment ions released from an *N*-glycopeptide containing the site N340-IgA1/N327-IgA2$_{(m1/n)}$. Oxonium marker ions confirming NeuAc O-acetylation (Acetyl$_1$NeuAc$_1$/316.1027 [M-H2O+H]$^+$, Acetyl$_1$NeuAc$_1$/334.1133 [M+H]$^+$ and HexNAc$_1$Hex$_1$Acetyl$_1$NeuAc$_1$/699.2454 [M+H]$^+$).

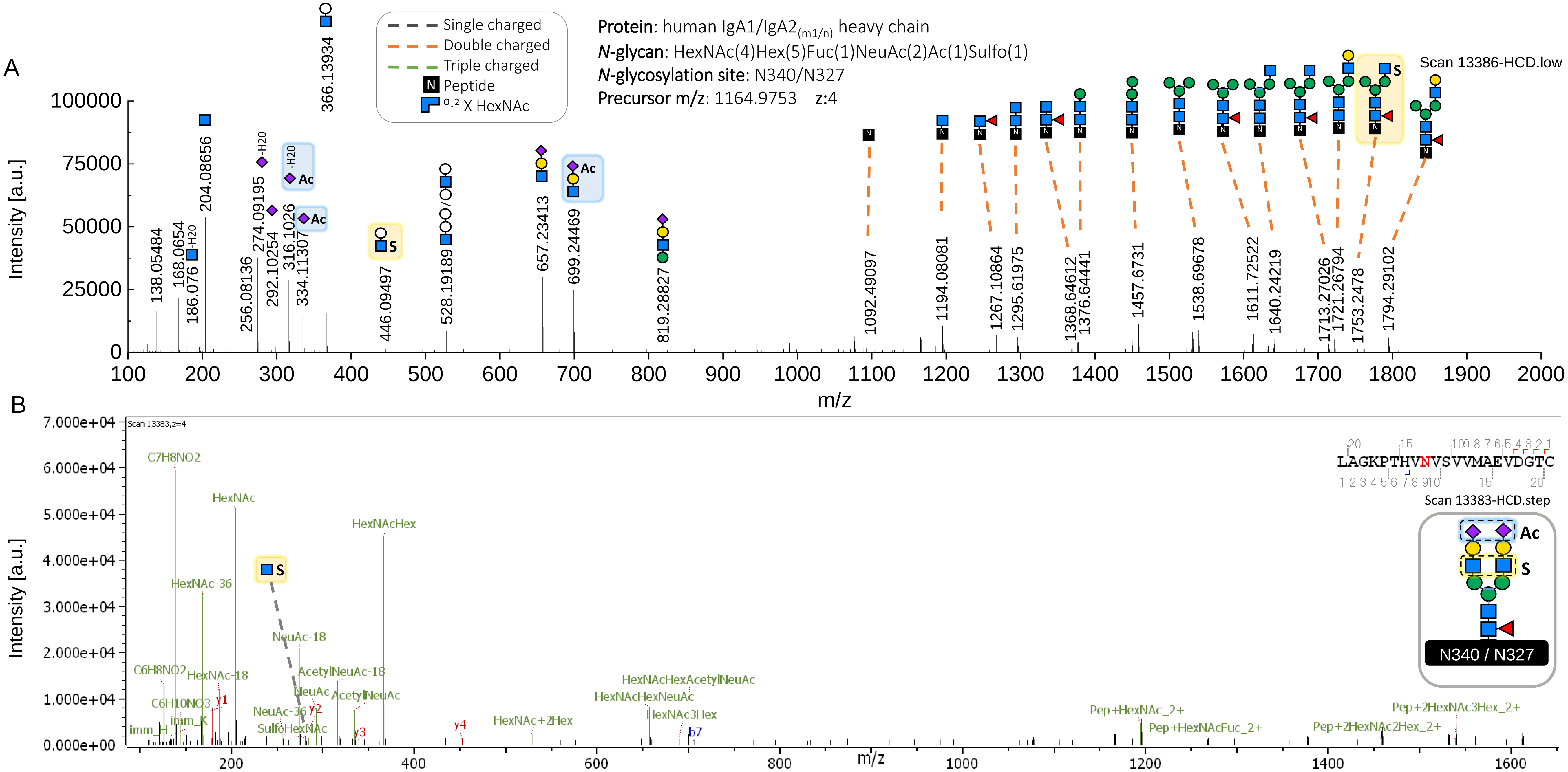


**Supplementary Figure 10. Glycopeptide containing NeuAc *O*-acetylation plus HexNAc sulfation in the *N*-glycan HexNAc(4)Hex(5)Fuc(1)NeuAc(2)Ac(1)Sulfo(1).** (A) Description of *N*-glycan structure through HCD.low spectrum shows B and Y ions providing structural evidence on NeuAc *O*-acetylation (highlighted in blue) and antenna sulfation (highlighted in yellow). (B) Byonic ™ annotation of the HCD.step spectrum identifies fragment ions released from an *N*-glycopeptide containing the site N340-IgA1/N327-IgA2$_{(m1/n)}$. The HCD.step spectrum also contains oxonium marker ions confirming HexNAc sulfation ($Sulfo_1HexNAc_1$/284.0431 $[M+H]^+$, highlighted in yellow) and NeuAc *O*-acetylation ($Acetyl_1NeuAc_1$/316.1027 $[M\text{-}H2O+H]^+$, $Acetyl_1NeuAc_1$/334.1133 $[M+H]^+$ and $HexNAc_1Hex_1Acetyl_1NeuAc_1$/699.2454 $[M+H]^+$).
